# Hummingbird iridescence: an unsuspected structural diversity influences colouration at multiple scales

**DOI:** 10.1101/699744

**Authors:** Hugo Gruson, Marianne Elias, Christine Andraud, Chakib Djediat, Serge Berthier, Claire Doutrelant, Doris Gomez

## Abstract

Iridescent colours are colours that change depending on the angle of illumination or observation. They are produced when light is reflected by multilayer structures or diffracted by gratings. While this phenomenon is well understood for simple optical systems, it remains unclear how complex biological structures interact with light to produce iridescence. There are very few comparative studies at interspecific level (often focusing on a single colour patch for each species), resulting in an underestimation of structure diversity. Using an interdisciplinary approach combining physics and biology, we here quantify the colour and structure of 36 hummingbirds species evenly distributed across the phylogeny. We explore at least 2 patches per species, which are assumed to be under different selective regimes. For each patch, we measure structural features (number of layers, layer width, irregularity, spacing, etc.) of the feathers at different scales using both optical and electronic microscopy and we measure colour using a novel approach we developed to encompass the full complexity of iridescence, including its angular dependency. We discover an unsuspected diversity of structures producing iridescence in hummingbirds. We also study the effect of several structural features on the colour of the resulting signal, using both an empirical and modelling approach. Our findings demonstrate the need to take into account multiple patches per species and suggest possible evolutionary pressures causing the evolutionary transitions from one melanosome type to another.

---

Hummingbirds are famous for their bright and shiny colours which change with the illumination or observation angle: a phenomenon known as iridescence. Iridescent colours are produced by the interaction of light with periodic nanometre-scale structures such as multilayers or diffraction gratings and are widespread among many taxa [1]. But few taxa display colours as bright and as saturated as the hummingbirds (Trochilidae family). Most hummingbird species harbour two visually distinct types of iridescent colour patches, as illustrated in fig. S1: *directional* patches, which are only visible at a very narrow angle range [2] and are often very bright and saturated, and *diffuse* patches, for which some colour is visible from any angle [2] and that are often not as bright as directional patches. Directional patches are often located on facial or ventral patches and thought to be involved in communication while diffuse patches are often located on dorsal patches and thought to be involved in camouflage [3]. Additionally, although all hummingbird species display some degree of iridescence, striking differences can be noticed between the various species and body patches in terms of brightness (describing how much light is reflected by the object), saturation (describing the colour “purity”) and directionality [4].

Yet, the structural bases of this intra-individual and interspecific diversity in colour have been poorly explored until now (but see Dorst [5]). In birds, multilayer structures responsible for iridescence are constituted of stacks of nanometre-scale melanin platelets or rods, sometimes hollow (i.e. with a central cavity filled with air) sometimes solid (i.e. entirely made of melanin), called melanosomes [6], included in a keratin matrix [7] (as illustrated in fig. 1). Although all of the 336 species in the family are iridescent [4], the multilayer structures of only 14 hummingbird species (represented on the hummingbird phylogeny in fig. S2) have been studied to this day [7–12]. These fourteen species all had hollow melanin platelets so this type of melanosome was assumed to be present in all hummingbird species [7]. However, studies in other families, such as starlings (Sturnidae), showed that multiple melanosome types can be present in the same family or even the same genus [7, 13], raising questions about the distribution of melanosome types and the evolution of iridescence in hummingbirds.

**Figure 1:**
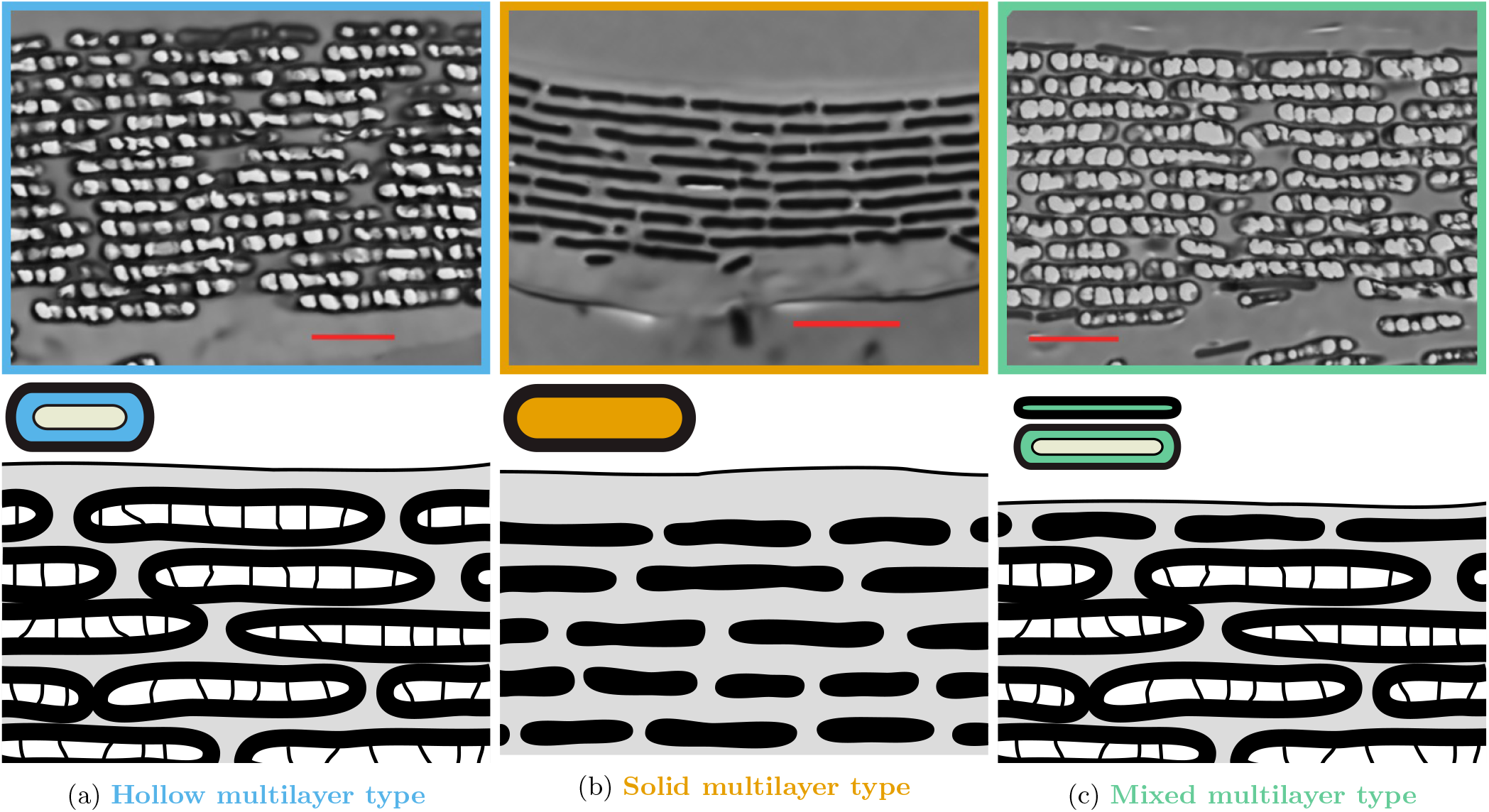
Examples of multilayer structures found in hummingbird barbules (top row) and their schematic representations (bottom row) adapted from Dürrer [7] with coloured symbols from Maia et al. [13]. The left panel shows hollow / air-filled platelets found in the breast of *Heliomaster furcifer*, which is the multilayer type that was known before for hummingbirds [7–10]. But we also discover two new types: the middle panel shows solid / melanin-filled platelets found in the back of *Aglaiocercus kingi* and the right panel shows a mixed multilayer structure with the outermost layer composed of solid / melanin-filled platelets and the rest of hollow / air-filled platelets, found in the throat of *Chrysolampis mosquitus*. The red bar represents 1 µm.

In this study, we aim at addressing three fundamental questions for the study of iridescence in hummingbirds but also in living organisms in general: 1) which type(s) of melanosome exist in hummingbirds and, if several exist, how are they distributed across the hummingbirds’ phylogeny? 2) do the different types result in different colour signals? 3) How do quantitative structural features (e.g. layer thickness, number of layers, etc.) influence the resulting colour?

To answer these questions, we adopted a mixed approach, using both empirical measurements on hummingbird iridescent feathers and transfer matrix optical simulations. We sampled one diffuse and one directional patch in 36 hummingbird species evenly distributed across the phylogeny (species position in the phylogeny shown in fig. S2).

Diffuse and directional patches are thought to be under different selection regimes and we accordingly formulate the following predictions: we predict that directional patches, which are often located on patches involved in communication, should reflect overall more light, and produce more saturated colours than diffuse patches, as these characteristics are often important in mate choice and quality advertising [14–17]. On the other hand, we predict that diffuse patches, which are often located on patches involved in camouflage should display a lower angle dependency of hue. Indeed, changes in colouration may cause “colour flashes” and alert a potential predator of the bird presence.

Additionally, hummingbirds present sickle-like shaped barbules [5, 7], illustrated in figs S3 and S4. We predict that this unusual shape may allow for a better interlocking of adjacent barbules and thus a higher spatial coherence, leading to a stronger interference pattern and ultimately brighter colours.

The detailed structural features of the multilayers for each patch were determined using Transmission Electron Microscopy (TEM) observations. For each patch, we also took colour measurements using a new method described in Gruson et al. [18] that allows the quantification of all iridescence characteristics, including angular dependency of hue and brightness. All analyses were performed by taking into account the phylogeny (comparative analyses), as to prevent pseudo-replication due to shared ancestry between species [19].

## Material and methods

### Colour measurements

We selected 36 species of hummingbirds evenly distributed across the phylogeny (see fig. S2; phylogeny data from Jetz et al. [20]). For each species (excepted for species that only had diffuse patches; see fig. 2), we sampled feathers on two patches, one diffuse (colour visible at many angles; often on dorsal patches) and one directional (colour visible at a small angle range; often on facial patches) from taxidermised specimens from the Muséum National d’Histoire Naturelle, in Paris. Feathers were carefully cut using surgical scissors and were only manipulated using tweezers, as to not remove or deposit any grease on the sample or modify barb arrangement.

**Figure 2:**
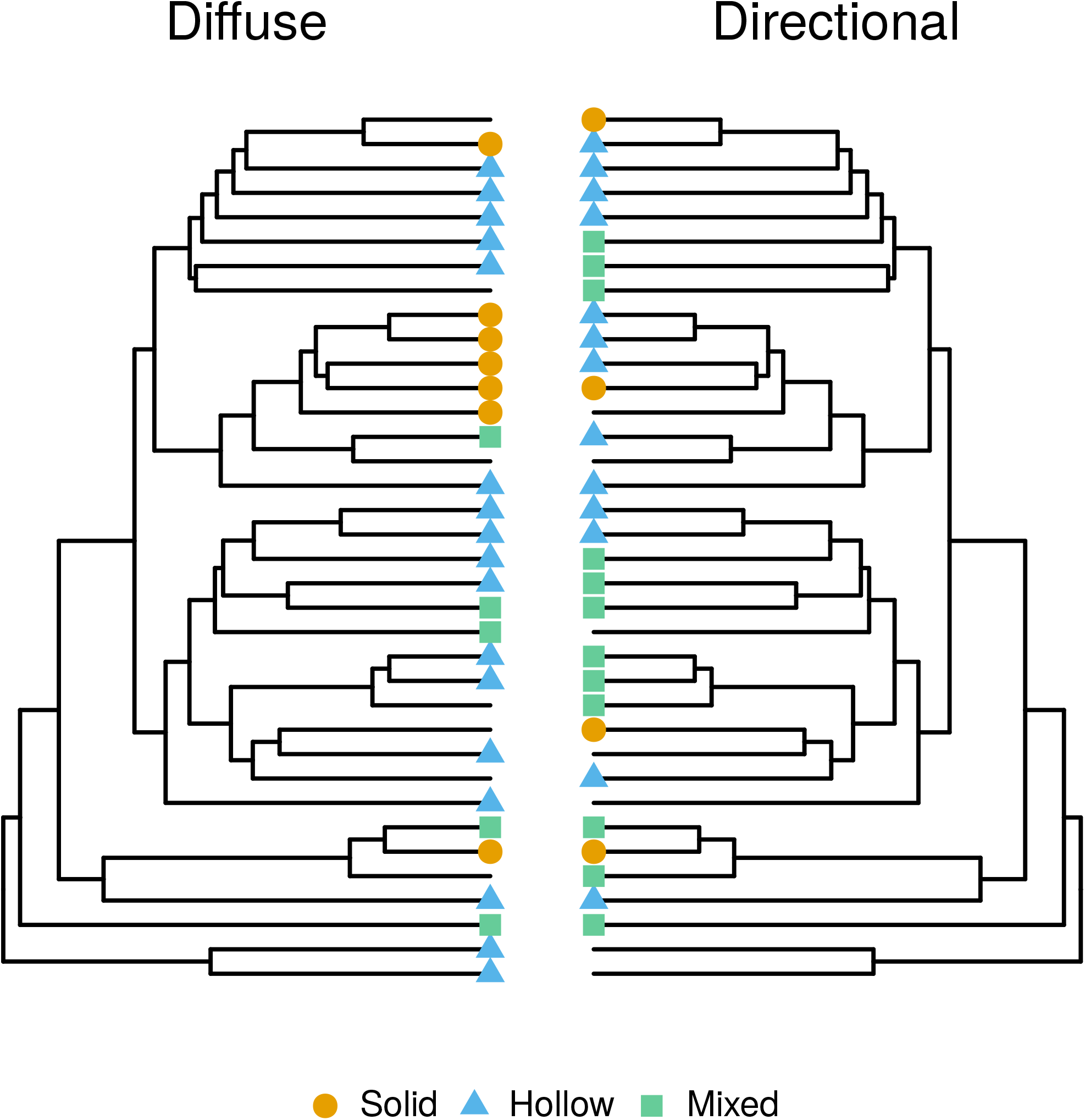
Some hummingbird species use different types of multilayer structures on different patches. For each species, we had one diffuse (left) and one directional (right) patch (diffuse and directional *sensu* Osorio and Ham [2]). Each tip is a species and tips are in the same order for both trees. Missing data are either species that do not have directional patches (e.g. *Patagona gigas* or species from the Hermit clade) or species that could not be measured due to technical issues. A more detailed version of this figure, with the type of multilayer for each patch and species names is available is SI (fig. S5).

Iridescence was quantified using the method published in Gruson et al. [18]. Briefly, we used a purpose-built goniometer to precisely quantify hue and brightness angular dependency in all directions. Using this method, brightness and its angular dependency can be summarised by two parameters: the maximum brightness *B*_max_ and the angular dependency of brightness *γ_B_* while hue and its angular dependency are defined by two parameters: the longest wave-length reflected *H*_max_ (reached when the observer and the incoming light are in the same direction) and the angular dependency of hue *γ_H_*. The saturation is expressed by the full width at half maximum (FWHM) of the spectra and does not change with the angle (low values of FWHM correspond to saturated colours). We recorded reflectance spectra with a 300 W Xenon lamp and an OceanOptics USB4000 spectrometer and two separate optical fibres for illumination and collection. All spectra were taken relative to a diffuse white spectralon standard (WS2 Avantes). Parameters were estimated using Bayesian non linear-regression with the brms R package [21, 22], which yielded slightly better results than non-linear least squares. All variables but the hue angular dependency *γ_H_* were repeatable between species, as reported in table S1. We also defined an additional variable called “overall reflectance” which takes into account both the specular and the diffuse reflectance of a sample and which is computed with the formula 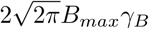 [18].

### Electronic and optical microscopy image acquisition and analysis

After colour measurement, we prepared feathers for observation with a Transmission Electron Microscope (TEM). Feathers were first dehydrated and then embedded in SPUR resin (detailed protocol in ESM). We used a Leica ultra-microtome to prepare 70 nm cross-sections of the barbules, where the multilayer structures responsible for iridescence are located [7]. We photographed the resulting cross-section with an optical microscope, which allowed us to measure the angle of the barbule (an measure of the barbule shape), the overlap between adjacent barbules, and the variance in the alignment of consecutive barbules (two measures of the interlocking between adjacent barbules). We then measured structural features at the scale of the multilayer such as the number of layers and their thickness using a TEM microscope (Hitachi HT-7700 TEM set at 60 keV).

Measurements on optical microscopy images were performed manually using the ImageJ computer software while TEM images were analysed using a custom python script, available in electronic supplementary materials (ESM), relying on the OpenCV python library [23, 24]. Briefly, we smoothed the grayscale images using Gaussian blur and a denoising algorithm. Resulting images were converted to binary black and white images using adaptive thresholding, then rotated using automatic contour detection, as to orientate the multilayer along the vertical direction. Finally, the number of transitions and the distance between them (layer thickness) in the rectangular function were determined for each row of the image matrix and the most common value was estimated using the mean of a fitted Gaussian function.

### Optical simulations

We used optical simulations to explore a wider combination of parameter values. The interest is twofold: 1) increase our limited sample size and 2) remove possible correlations (possibly due to evolutionary constraints) between structural parameters.

We used the EMpy python library [24, 25], which implements the transfer matrix method described in Yeh [26] to simulate the reflected specular spectrum of a multilayer structure. The script used for the simulations is also provided in ESM.

Because of the large array of parameters influencing the resulting reflectance spectrum (complex optical index of each layer, layer thicknesses, angle of the incoming light ray, number of layers), it was not possible to systematically study the effect of each parameter. To overcome this issue, we ran 500 iterations of Monte Carlo simulations, for each multilayer type, with structural parameters randomly drawn from an interval of biologically relevant values. This interval was determined from the TEM images (95 % variation interval for each parameter, irrespectively of the multilayer type). We had several images for each species and patch combination, which allowed us to ensure that all estimated structural variables were repeatable (table S1).

Because there is no disorder in the layer alignment, the brightness in the simulations corresponds to the overall reflectance (diffuse + specular reflectance) in the empirical measurements (computed with the formula 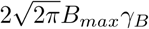).

The results are presented in SI with only the mean and the standard deviation of the parameter influence on the response variable, as appropriate for simulations (because the possibly infinite sample size allows for arbitrary low p-values). Additionally, significance of the effect of a given parameter for a sample size of 72, to match the sample size of empirical data, is shown in table 1, using Cohen’s d effect size index.

**Table 1:** Predicted correlations between colour variables and structural parameters and the outcome from comparative analyses and simulations for these correlations. The correlations can be due to either the optics governing iridescence and to evolution. As mentioned in the methods, it is possible to get an arbitrary low p-value in simulations by increasing the sample size. To prevent this issue and to be able to compare empirical and simulations results, we chose the same sample size for both (72) and counted a result as significant only when *p <* 0.05 (for simulations) or when the credibility interval did not include 0 (for empirical data). Some correlations could not be tested in the simulations and are marked as NA in the table. If results from the empirical data and the simulation output the same result, it is likely due to the optics governing iridescence but in case of mismatch, it reveals the influence of evolutionary constraints.

|  | Variable 1 | Variable 2 | Empirical | Simulations |
| --- | --- | --- | --- | --- |
| Colour/structure | Hue at coincident geometry ($H_{max}$) | Layer thickness | yes | yes |
| | Hue at coincident geometry ($H_{max}$) | Multilayer type | yes | no |
| | Overall reflectance ($2\sqrt{2\pi}\gamma_B B_{max}$) | Number of layers | no | no |
| | Overall reflectance ($2\sqrt{2\pi}\gamma_B B_{max}$) | Multilayer type | yes | yes |
| | Overall reflectance ($2\sqrt{2\pi}\gamma_B B_{max}$) | Barbule shape | yes | NA |
| | Directionality ($1/\gamma_B$) | Barbule shape | yes | NA |
| | Directionality ($1/\gamma_B$) | Multilayer type | yes | NA |
|  | Desaturation (FWHM) | Multilayer type | yes | yes |
|  | Desaturation (FWHM) | Barbule shape | no | NA |
|  | Desaturation (FWHM) | Layer thickness variance | no | NA |
|  | Desaturation (FWHM) | Number of layers | no | yes |
|  | Desaturation (FWHM) | Layer thickness | NA | no |
|  | Hue shift | Multilayer type | NA | yes |
| Colour/Colour | Hue at coincident geometry ($H_{max}$) | Overall reflectance ($2\sqrt{2\pi}\gamma_B B_{max}$) | yes | no |
| | Overall reflectance ($2\sqrt{2\pi}\gamma_B B_{max}$) | Directionality ($1/\gamma_B$) | yes | NA |
| | Overall reflectance ($2\sqrt{2\pi}\gamma_B B_{max}$) | Desaturation (FWHM) | yes | yes |
| | Desaturation (FWHM) | Directionality ($1/\gamma_B$) | no | NA |
| | Hue at coincident geometry ($H_{max}$) | Desaturation (FWHM) | yes | yes |

We also analysed the resulting spectra as seen by the hummingbirds using Stoddard and Prum [27] model, implemented in Maia et al. [28?]. The gamut of each multilayer type was computed as the volume of the convex hull of the set of points in the tetrahedron representing bird colour space, as in [27].

### Predictions

We can formulate a set of predictions for correlations between colour variables and structural parameters, as well as among colour variables, based on two factors: (i) predictions informed by optical theory and the laws of interferences from multilayers (ii) predictions informed by previous research on colours as a communication channel in animals.

In particular, based on the equation computing the wavelengths at which reflected light rays interfere constructively the most, *mH_max_* = 2(*n*_1_*e*_1_ +*n*_2_*e*_2_), we predict that hue (*H_max_*) and the angular dependency of hue (*γ_H_*) should depend on layer thickness (*e*_1_; *e*_2_) and chemical composition (*n*_1_; *n*_2_), as well as interference order (*m*). The angular dependency of brightness *γ_B_* should only depend on the misalignment between consecutive layers or multilayer, because a perfectly aligned multilayer should reflect all light in a single direction (*γ_B_* = 0), as detailed in Gruson et al. [18]. Total reflectance 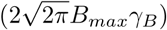 is expected to be positively correlated to the number of layers (because more light is reflected and more rays interfere), the chemical composition (melanin layers absorb more light) and the spatial coherence of adjacent multilayers (influenced by the barbule shape and the amount of overlap between adjacent barbules). Finally, saturation should depend on the variability in layer thicknesses (because it produces a mix of wavelengths that constructively interfere), the misalignment of consecutive layers, as well as the number of layers and the chemical content (because selective absorbance of some wavelengths would increase saturation).

We do not study maximum brightness *B_max_* separately as we have shown before that it is strongly correlated with *γ_B_* because of structural reasons (Gruson et al. 18; illustrated also in fig. S9).

Additional predictions are due to the putative function of iridescent colours in hummingbirds: colour on directional patches should be highly saturated and reflect overall more light than on diffuse patches, as directional patches are thought to be involved in communication and high brightness and saturation are common quality indicators [14–17]. In other words, we predict a negative correlation between *γ_B_* and the FWHM (measure of desaturation, opposite of saturation) as well as overall reflectance.

### Correlations between structure and colour using phylogenetic comparative analyses

The different multilayer structures studied in this article are not independent samples from a statistical point of view. Indeed, all samples come from species that share a common evolutionary history. This shared history, represented by species phylogeny, must be taken into account using phylogenetic comparative analyses [19]. However, classic phylogenetic comparative methods do not consider multiple data points per species. Since we measured two patches per species, we used the Bayesian framework implemented in the R package `MCMCglmm` [29, 30], which allows analysing such data (see Delhey et al. [31] for another example). Multiple studies have shown that results from comparative analyses are more reliable when performed on multiple trees drawn from the posterior distribution instead of a consensus tree [32, 33]. To account for such phylogenetic uncertainty, we ran models using the `mulTree R` package [30, 34] on a set of 100 trees dowloaded from birdtree.org [20]. Each model was tested with 3 independent MCMC chains, with 200 000 iterations each, including a 1000 burn-in and a thinning factor of 10 to reduce auto-correlation and memory consumption of the program. Convergence was assessed both visually and using the Gelman-Rubin index. Levels of a factor were deemed significantly different when the estimate of one did not overlap with the credibility interval of the other.

Phylogenetic signal for the type of multilayer on the throat and the back was computed using the *δ* Bayesian approach for discrete characters described in Borges et al. [35]. Larger values of *δ* express a higher level stronger phylogenetic signal, i.e. a stronger influence of the evolutionary history on the observed trait values. *δ* values can be arbitrarily large and significance is evaluated by bootstrapping after shuffling the trait value on the phylogeny.

## Results

### Correlations between colour variables in hummingbird iridescent feathers

Preliminary study of correlation between colour parameters, without investigating yet the underlying structural variable, reveals a positive correlation between maximum brightness *B_max_* and saturation table S2. Directional patches (low *γ_B_*) also tend to reflect overall more light than diffuse colours (table S3). On the other hand and contrary to our predictions, we find no correlation between *γ_B_* (related to directionality) and saturation (table S2). We also find that long wavelength hues (i.e. red colours) are associated with brighter (table S2) but less saturated colours (table S3).

### Iridescence in hummingbirds is produced by several different multilayer types

Observations of barbule cross-sections with a Transmission Electron Microscope (TEM) confirm that some hummingbird multilayers contain only hollow melanosomes (left panel in fig. 1). But we also discover that some species have multilayers with solid melanosomes (central panel in fig. 1). Additionally, we find a highly unusual multilayer structure in some species, where the outermost layer is composed of solid melanosomes while the rest of the multilayer is composed of hollow melanosomes (right panel in fig. 1). We refer to this multilayer structure type as the *mixed* multilayer type in the rest of this article. Lastly and importantly, our observations show for the first time that a single hummingbird species can have different multilayer types depending on the patch location on the body as shown in fig. 2 and fig. S5. The thickness of the melanin layer is very similar between hollow and solid melanosomes (fig. S6). However, because solid melanosomes contain only one layer of melanin (versus two layers of melanin surrounding one layer of air for hollow melanosomes), they are overall much smaller than hollow melanosomes. Hollow melanosome thicknesses range from 130 nm to 228 nm with the air void filling on average 44 % of the total thickness, while solid melanosomes measure between 29 nm and 80 nm. The total number of melanin layers (2 per hollow melanosomes vs 1 per solid melanosomes) does not significantly differ between the multilayer types (fig. S6). More detailed data, including variation intervals, relative standard deviations and repeatabilities for each parameter, is presented in table S1 and fig. S6.

### Location on the bird body and optical effects of the different types of multilayers

We find that diffuse patches contain multilayers with only hollow melanosomes more often than directional patches. At the same time, directional patches contain mixed multilayers more often than diffuse patches (*χ*^2^(2) = 6.8138, *p* = 0.033; fig. S10).

There is also a strong phylogenetic signal for the multilayer type on the back (*δ* = 11.03, *p* = 0.008) but not on the throat (*δ* = 1.37, *p* = 0.067).

Using phylogenetic comparative analyses, we also find that multilayer structures with only hollow melanosomes reflect overall more light (diffuse + specular reflectance; table S8) but less saturated colours (i.e. larger FWHM, table S11) than structures with solid melanosomes. Mixed multilayers have intermediate values compared to solid and hollow multilayer types for both brightness and saturation. This result is confirmed by transfer matrix simulations, which allow us to explore a much wider range of parameters and ensure this pattern is not caused by a confounding effect (fig. S12 and tables S4 and S5).

The different multilayer types also produce different hues, with the mixed type producing the largest diversity of hues in the bird visual space, using simulations based on biologically relevant layer sizes (fig. S13).

Finally, the different multilayer structures also differed in their level of iridescence, i.e. how much hue shifts with a change in the angle of illumination or observation, with the hollow type having a larger shift in hue than the solid and the mixed types in the simulations (fig. S11 and table S7). This could not be verified on empirical data with hummingbird feathers as this variable was not repeatable (table S1).

### Optical effects of structural features

At the multilayer level, the number of layers has no effect on overall reflectance in phylogenetic comparative analyses based on empirical data from hummingbird feathers (table S8). Simulations similarly reveal a very weak correlation between the number of layers and brightness (fig. S12 and table S4). On the other hand, a larger number of layers did increase saturation for both hollow and solid multilayer types but not for the mixed type in the simulations (fig. S12 and table S5). Variability in the thickness of melanin, keratin or air layers of a given multilayer did not seem to significantly impact the saturation of the resulting signal (table S5).

We show that hue at a given angle configuration (*H_max_*) depends on the thickness of the layers, no matter their chemical composition (air, keratin or melanin), in simulations (table S6) but we only find a significant effect of the thickness of the melanin layer in empirical data (table S10). However, we also find that thicknesses of melanin, keratin and air layer within a given multilayer structure are strongly correlated, as shown in fig. 3, which might hinder our analysis on empirical data. This correlation is not simply due to phylogenetic inertia and the shared history between species as it remains significant even after taking into account the species phylogenetic relationships (tables S12 to S14). Additionally, the confidence interval of the slope of the correlation between the optical thicknesses (thickness times optical index) of the consecutive layers is often close to 1 but does not contain 1, as shown in fig. 3.

**Figure 3:**
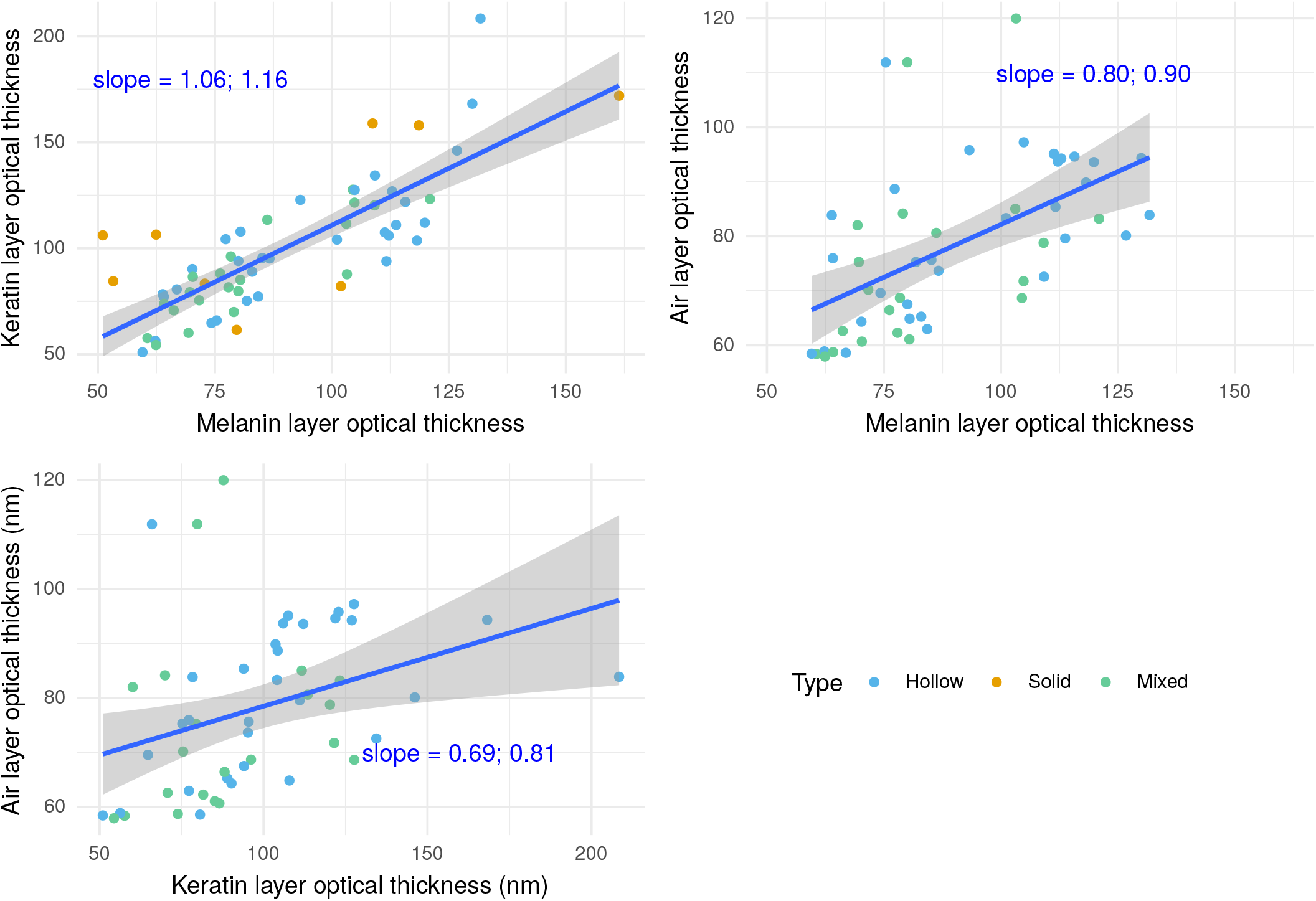
Optical thickness of melanin, keratin and air layers are correlated in hummingbird multilayer structures. Each dot is a multilayer from a given species/patch combination. Correlations are tested by linear models (blue lines on the present figure. Confidence interval of the slope is in blue as well.) which do not take into account species relatedness, and by comparative phylogenetic analyses using MCMCglmm (tables S12 to S14). There is no data for air layer thickness in solid multilayer types because they do not contain any air.

At the barbule level, we find that barbules with a sickle shape (i.e. with a smaller barbule shape angle, as shown in fig. S4) produce colours that reflect overall more light (taking into account both diffuse and specular reflection), as illustrated in fig. 4b and table S8. Additionally, in agreement with our predictions, we show that barbules with a sickle shape also produce more directional and more saturated colours (fig. 4a and tables S9 and S11). Conversely, variability in barbule alignment from the same barb also produces less saturated colours (table S11).

**Figure 4:**
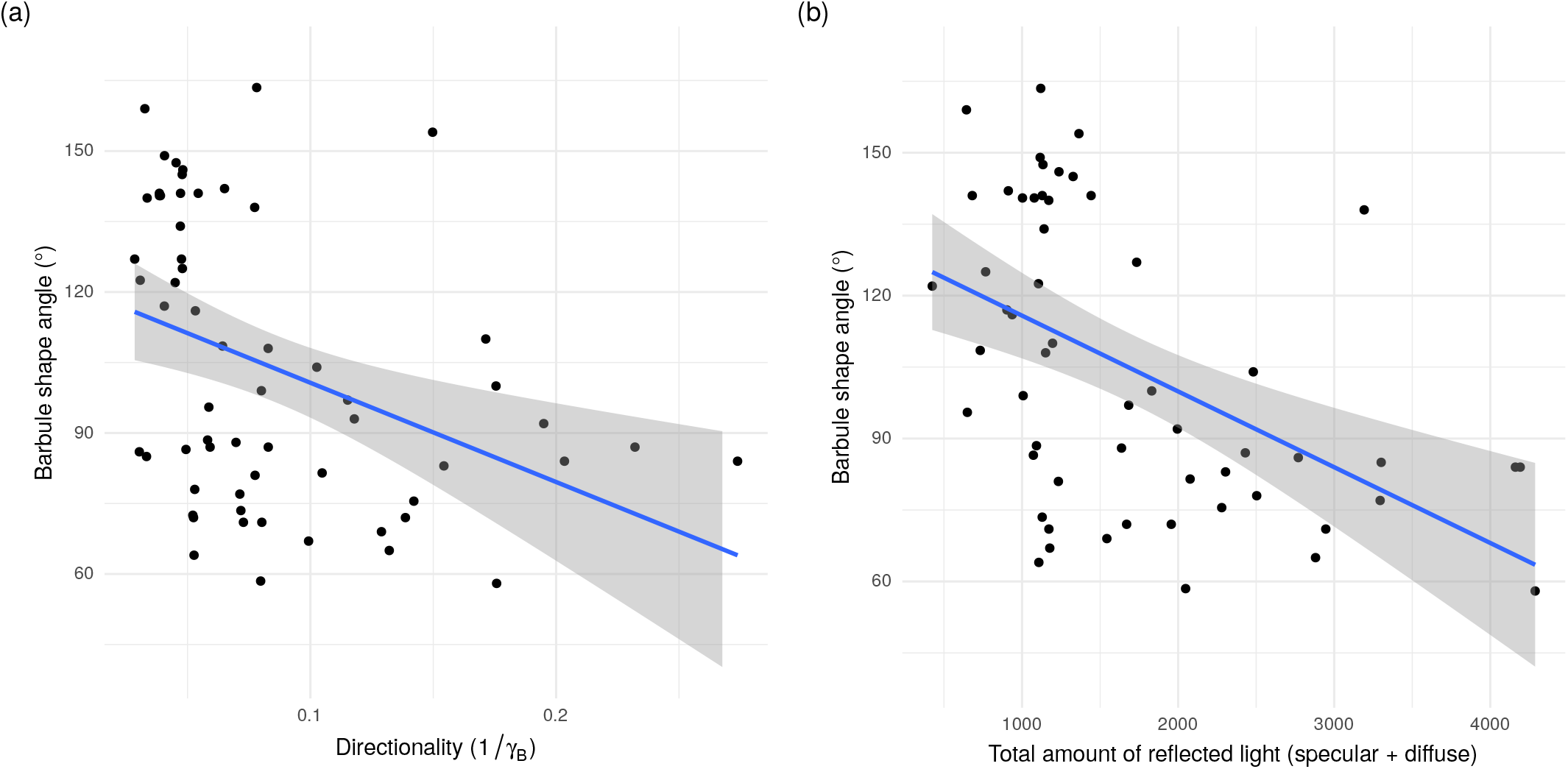
Link between the barbule shape angle and (a) the directionality 1*/γ_B_* or (b) the total amount of reflected light (specular + diffuse reflected light; computed with the formula 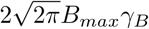 from Gruson et al. [18]). Barbules with a sickle shape (lower barbule shape angle) produce on average colours which are more directional (lower value of*γ_B_*) and reflect overall more light (taking into account both diffuse and specular reflection). Regression line in blue and the related 95 % confidence interval in grey are fitted by a linear model and only have an illustrative function. Phylogenetic comparative analyses which take into account phylogenetic inertia are presented in table S8.

## Discussion

### Correlations and general characteristics of hummingbird iridescent colours

We find many correlations between descriptors of iridescent colours in hummingbirds. In particular, saturation was negatively correlated with hue (table S2), as expected for interferences from a multilayer structure. For long wavelength colours, a wider range of wavelengths will indeed (partially) constructively interfere and contribute to the resulting signal, thereby producing less saturated colours. Our framework did not allow us to discriminate whether evolutionary constraints could also play an additional role in the correlation (i.e. is there a selective pressure for blue colours to be more saturated than red colours in hummingbirds?)

We nonetheless found additional correlations that are not explained by the physical nature of hummingbird colours. For example and in accordance with our prediction, we found a positive correlation between saturation and total reflectance, as could be expected from patches involved in quality advertising and mate choice [14–17].

Finally, we showed a correlation positive correlation between hue and overall brightness, meaning that red colours are on average brighter than blue colours. Two non-mutually exclusive hypotheses can explain this pattern: (i) red colours are often used for communication in hummingbirds due to a pre-existing sensory bias [36] and communication is often associated with brighter colours, or (ii) blue colours are not as bright because melanin and keratin absorb more in short wavelengths than in long wavelengths [37, 38].

### Hummingbirds display an unsuspected multilayer diversity

In this study, we discover that hummingbirds do not only have hollow (air-filled) melanosomes but also solid (melanin-filled) melanosomes. They also sometimes combine both types into a very unusual multilayer structure that has never been described in any other family, where the outermost layer is formed by solid melanosomes while the following layers contain hollow melanosomes.

We also discovered that a single species can use different types of multilayer structures at different patch locations on its body (fig. 2 and fig. S5). This means that the type of multilayer found on one patch is not representative of the multilayer type found on all patches for a given species. This finding calls for more careful investigation into the results of previous comparative analyses of bird melanosomes and iridescent colours, as most of them have observed only one patch per species [7, 13].

We also show that the different types of multilayers are not randomly distributed on the bird’s body: diffuse patches contained multilayers composed exclusively of hollow or solid melanosomes more often than directional patches. On the other hand directional patches contained mixed multilayers more often than diffuse patches (fig. 2 and fig. S10). We find a strong phylogenetic signal for the type of multilayer structure on the back but not on the throat, suggesting that the distribution of the multilayer type is mainly due to the phylogeny on the back but likely more strongly influenced by additional selective pressures on the throat.

This suggests that the different multilayer types produce different kinds of colours that are selected in different contexts: mixed types may produce colours that are generally more efficient for communication while hollow or solid types produce colours more efficient for camouflage.

### Different multilayer types produce different colours

For hue, and in conformity with our prediction that diffuse patches should contain multilayer structures that minimise the angle dependency of hue, we found that diffuse patches contained the solid multilayer type more often than directional patches, which leads to a lower hue shift in simulations (fig. S11 and table S7). We could not verify this prediction with empirical data as *γ_H_* was too similar across species to yield repeatable measurements. This lower hue shift could reduce colour flashes that may alert a potential predator of the presence of the bird. On the other hand, diffuse patches have most commonly hollow melanosomes, which can lead to the highest hue shift (fig. S11 and table S7).

This partial mismatch with our prediction could be explained by the findings of Kjernsmo et al. [39], where the authors found that iridescence could improve camouflage by impairing predators’ ability to discern target shape.Alternatively, the difference in hue shift among the different multilayer types could be low enough to not be under strong selective pressure.

We also found with simulations that the mixed multilayer type can produce the highest diversity of hues (fig. S13), while the solid type has the lowest diversity. It does however seem that the full range of possible hues is not explored in hummingbirds. This is probably in part due to our non-exhaustive sampling of hummingbird species but also likely reflects evolutionary constraints, either on the structures themselves or on the resulting colour [3].

For brightness, previous studies predicted based on optical theory that hollow melanosomes should produce brighter colours than solid melanosomes [40, 41]. The simulations in the present study confirm that multilayers with hollow melanosomes reflect more light overall (specular + diffuse reflectance) than multilayers with solid melanosomes. Mixed multilayer types have intermediate values between multilayers with only solid or hollow melanosomes (table S8). However, the multilayer type is likely to have a minimal effect on effective brightness at a given angle. The bright colours of hummingbirds are indeed not caused by an increase of the total amount of reflected light but rather by a very high directionality of the reflected signal, meaning that all reflected light is focused within a narrow angular sector, as found by Osorio and Ham [2], Gruson et al. [18] and shown for our study in fig. S9.

On the other hand, we find that, in both empirical data and simulations, hollow multilayers produce less saturated colours than multilayer structures with solid melanosomes or mixed multilayers, as shown in table S11 and fig. S12 and table S5 respectively. However, the mixed multilayer type had the highest interaction value with the number of layers (fig. S12 and table S5). This means that mixed multilayers have the highest potential to create highly saturated colours when composed of a large number of layers, which could explain that they were positively selected in directional patches.

Our results describing the influence of the multilayer type on brightness and saturation are also in line with the detailed study of Giraldo et al. [11] on the throat feathers of Anna’s hummingbird (*Calypte anna*): using optical simulations, they found that the exclusion of the thinner outermost layer in their simulations produced brighter but less saturated colours.

From a macro-evolutionary point of view, the evolution of new types of multilayer structures might also be responsible for the rapid diversification rate of hummingbirds [42], playing the role of key innovations that allow them to quickly fill up previously unexplored regions of the phenotypic space (more specifically, new hues and more saturated colours, as mentioned above), as was previously described in iridescent starlings by Maia et al. [13].

### Multilayer structures in hummingbirds are not very regular but often close to ideality

We found a very high intra-multilayer variability for the structural characteristics of melanosomes, as expressed by the high relative standard deviation values reported in table S1. These values are close to previous values reported in the literature for hummingbird multilayers [12], and they likely reflect actual biological variability rather than measurement uncertainty. For example, Greenewalt et al. [8] found that layer thickness generally varied between 20-30 % within species. The thickness we measured for hollow melanosomes (95 % variation interval = 130 nm to 231 nm) was also well within the range of what was estimated in the past on a smaller species sample (100 nm to 220 nm, with a mode of 150 nm for Greenewalt et al. [8] and between 200 nm and 250 nm for Dorst [5] with a photonic microscope).

We also observe strong correlations between the thickness values of the different layers, as shown in fig. 3. In other words, melanosomes with a thicker layer of melanin were also spaced by thicker layers of keratin. This correlation could explain the above mentioned fact that realised hues are much less diverse than possible theoretical hues for each multilayer type using simulations with biologically relevant ranges for layer thicknesses. This correlation is not caused by the phylogenetic relationships between species and remains significant even when the phylogeny is taken into account (tables S12 to S14). This suggests the existence of selective pressures that maintain this correlation. This may be due to selection for ideal multilayers, where the optical thickness (defined as optical index times layer thickness *n_i_e_i_*) of the successive layers is constant [43]. This hypothesis is supported by the fact that slopes of the correlations between optical thicknesses of successive layers are close to 1 (fig. 3). Ideal multilayers are found in many organisms such as the butterfly *Chrysiridia rhipheus* [44], *Sapphirina* copepods [45] or the Japanese jewel beetle *Chrysochroa fulgidissima* [46] where they are thought to be selected because they produce brighter, more saturated colours [43].

Because the wavelength (i.e. hue) reflected by a multilayer depends on the thickness of the layers, variability in thickness may produce a mix of numerous wavelengths, and thus less saturated colours. This prediction is however not supported by our results (table S11), which suggests that saturation is not significantly explained by variability in layer thickness.

### Multilayer types correlate with other structural features that enhance conspicuousness

We found little or no effect of the number of layers on brightness, in both simulations (table S4) and comparative analyses (table S8). This result is in agreement with what Eliason et al. [47] found in melanosome rods from dabbling ducks and could be partly explained by the fact that brightness quickly reaches a plateau when the number of layers increases [43, 48, 49]. Indeed, multilayer theory predicts that brightness increases exponentially towards its maximum with the number of layers [26]. For example, Giraldo et al. [11] found that 10 layers created a spectrum that was close to saturation in their modelling investigation of the pink throat feathers from Anna’s hummingbird (*Calypte anna*). Similarly, Stavenga et al. [50] reported that 5 layers were sufficient to reach the maximum brightness in their study on magpie (*Pica pica*) hollow melanin rods while Berthier et al. [51] wrote that less than 10 layers achieved maximum brightness in an ideal chitin-air multilayer.

However, the multilayers found in hummingbirds have at least 5 layers, with a median of 12 layers and a maximal number of layers sometimes over 25, well beyond the theoretical number of layers needed to reach maximum reflectance. Indeed, an increasing number of layers did increase saturation (i.e. decrease FWHM) in simulations (fig. S12 and table S5). This result suggests an explanation for the unusually high number of layers in hummingbird feathers, and especially on directional patches. As we mentioned earlier, more saturated colours are often positively selected in the context of communication. Selection for higher saturation could then be the driving force for the evolution towards a higher number of layers in patches used in communication, even though brightness does not significantly increase.

We found that the sickle shape of hummingbird barbules is correlated with more reflective (table S8, more saturated (table S11) and more directional colours (table S9). It remains unclear whether barbule shape has an optical role at the level of the barbule itself as is the case for example in the triangular barbules from the breast of the bird of paradise *Parotia lawesi* [52, 53]. However, it is unlikely that this shape contributes directly to the interference pattern because the position of the lower part of the “sickle” (also called *velum*) does not reflect light rays in the same direction as the upper part (also called *speculum*) [11]. However, multiple studies have suggested that barbule organisation nonetheless influences the resulting signal (see for example Schmidt and Ruska [9] on the hummingbird *Heliangelus strophianus* or Dürrer and Villiger [54] on the golden cuckoo *Chrysococcyx cupreus*). Indeed, more packed barbules produce brighter colours because the light reflected from each multilayer also interferes at the level of the barb or even the whole feather. This peculiar sickle shape could then have been selected because it allows for a better interlocking of adjacent barbules, leading to a greater spatial coherence across scales, causing signals reflected by each barbule to interfere more constructively and ultimately produce brighter, more saturated colours. This stronger interlocking could also have an effect on processes other than colour generation, such as producing more waterproof feathers, or increasing lift during flight by limiting air gaps between barbules, which may be especially important for the stationary flight of hummingbirds [55]. Dorst [5] suggests that mechanics (for flight) and optics (for colour) benefit from the same modification and selection likely acts on both jointly.

There are other structural parameters we could not measure with our present experimental setup but that could influence the resulting colour; namely the angle between the barbules and the parent barb in the plane of the feather (named ‘barbular angle’ in Greenewalt [56] and represented in fig. S4), the angle between barbules and the barb axis in a cross section of the barb (named ‘vanular angle’ in Greenewalt [56] and represented in fig. S4). However, Dorst [5] found no effect of the barbular angle on the visual appearance of the feathers in his investigation of 15 hummingbird species.

## Conclusion

The present study sheds a new light on the evolution of iridescence, in hummingbirds, and more generally in all other organisms, with several major findings: (i) hummingbirds display much more diverse multilayer structures than previously expected, with even a type of structure unknown thus far, (ii) a single species may display multiple types of multilayer at different location on its body, and (iii) structural features at both the level of the multilayer and the level of the whole feather interact in the production of iridescent colours.

## Acknowledgements

We thank the Museum National d’Histoire Naturelle (Paris) for allowing us to sample hummingbird feathers for TEM observations and iridescence measurement. In particular, we are grateful to the museum curators Patrick Boussès, Jérôme Fuchs and Anne Previato for their help. We also thank Nicolas Moreau and Colin Durand for their assistance with photonic microscopy photographs.

## Authors contributions

HG, ME and DG designed the study and discussed it with CDo. HG and CDj performed TEM observations. HG, SB and CA discussed colour measurement protocol and transfer matrix simulations. HG performed colour measurements, structure measurements on TEM images, and simulations. DG analysed structures on photonic images. HG, DG, and ME worked on comparative analyses. All authors contributed to the final version of this manuscript.

---

**Supplementary figure 1:**
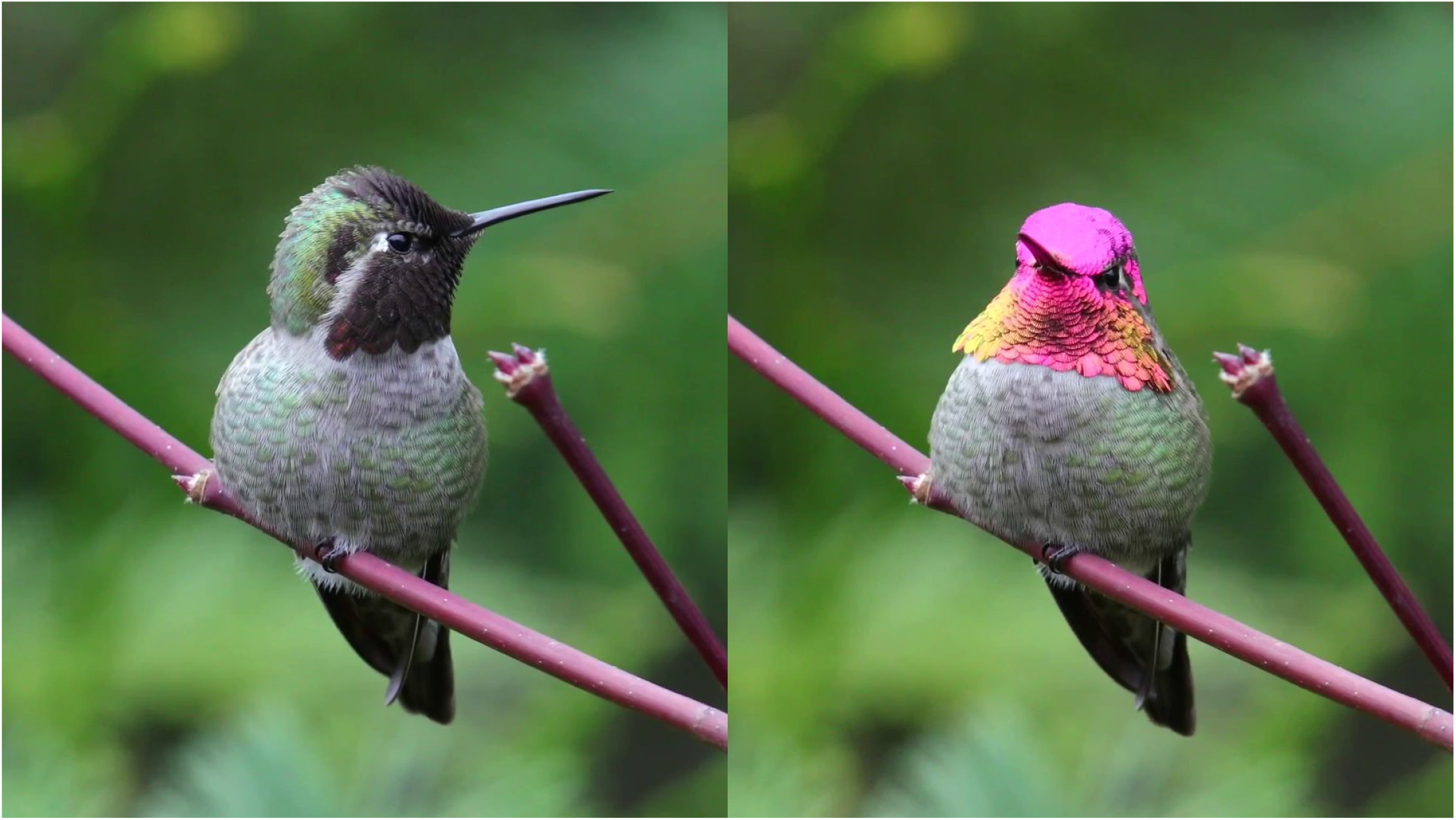
Picture of the same *Calypte anna* individual at different angles. Modified from a video taken by Mick Thompson (https://www.flickr.com/photos/mickthompson/27991602299/), CC-BY-NC, special authorisation to use it in this article. The directional throat and crown patches contrast with the diffuse greenish belly patch.

**Supplementary figure 2:**
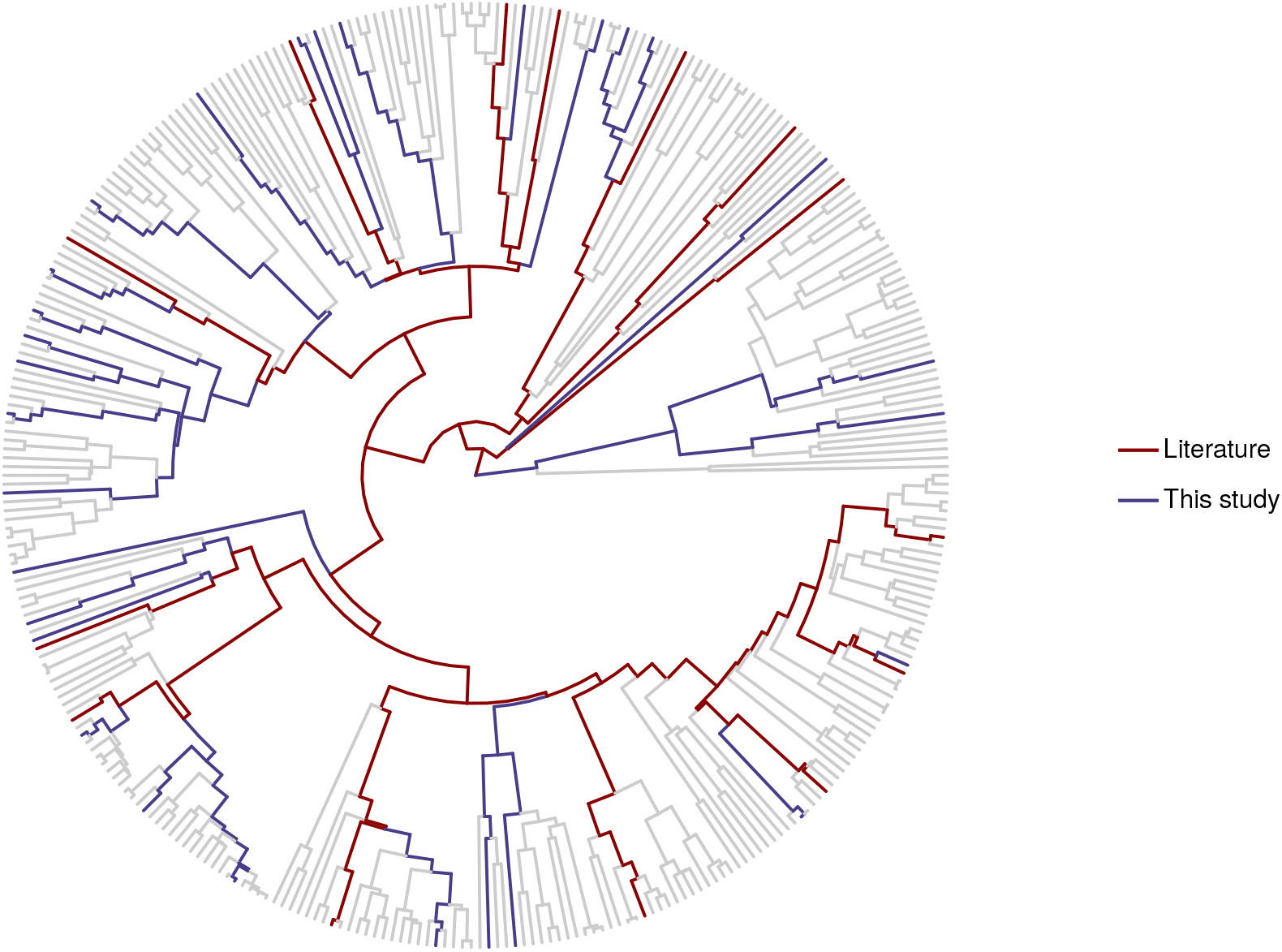
Consensus phylogeny of the hummingbirds reconstructed using the Maximum Clade Credibility tree from a distribution of 4999 trees downloaded from http://birdtree.org [20]. The red lineages show the 14 species whose structures had been previously studied in the literature [7–10, 12] while blue lineages show species we studied for the first time in this study.

**Supplementary figure 3:**
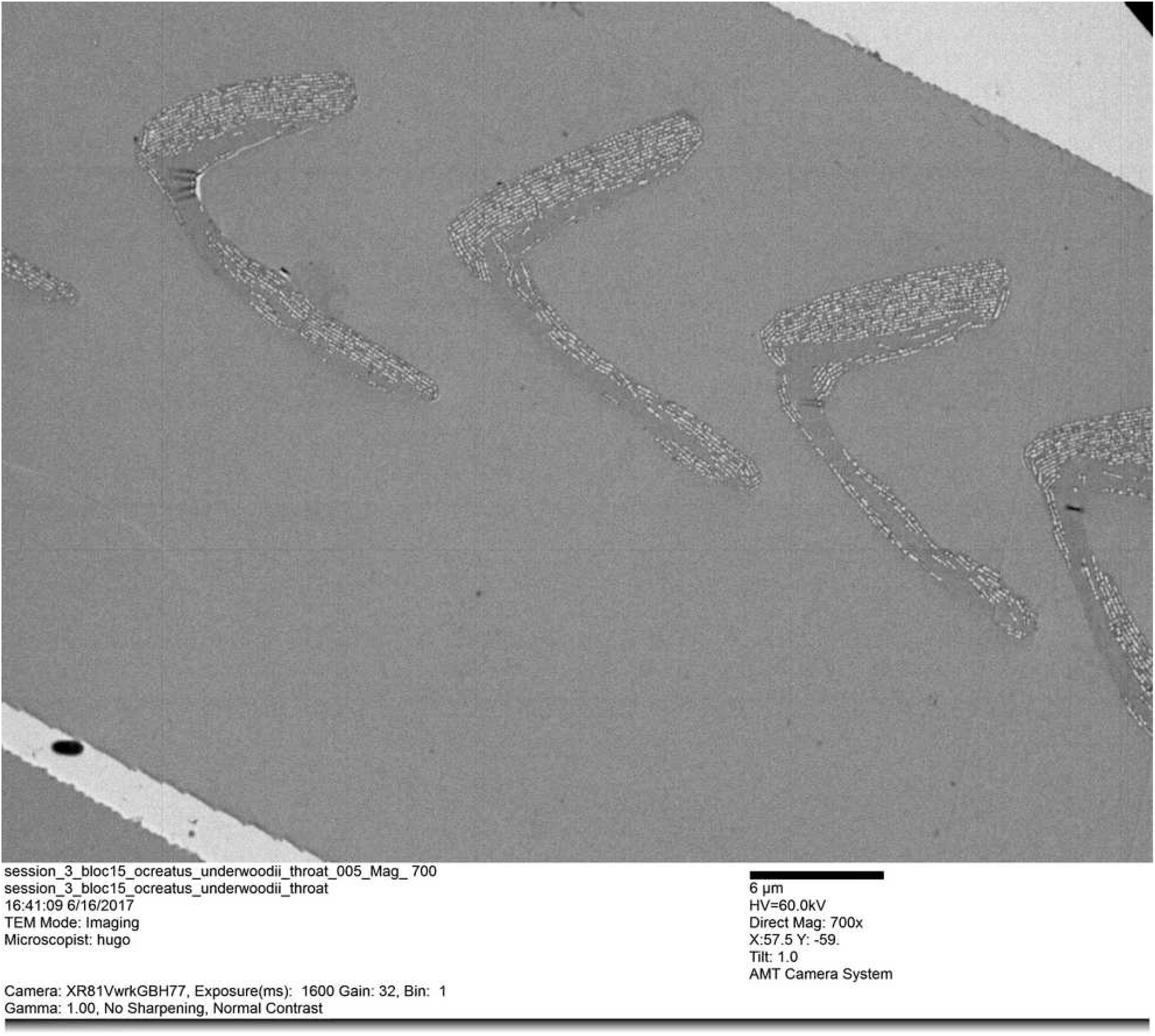
Unedited TEM photograph of the cross section of 5 consecutive barbules from the same barb of the throat of *Ocreatus underwoodi*.

**Supplementary figure 4:**
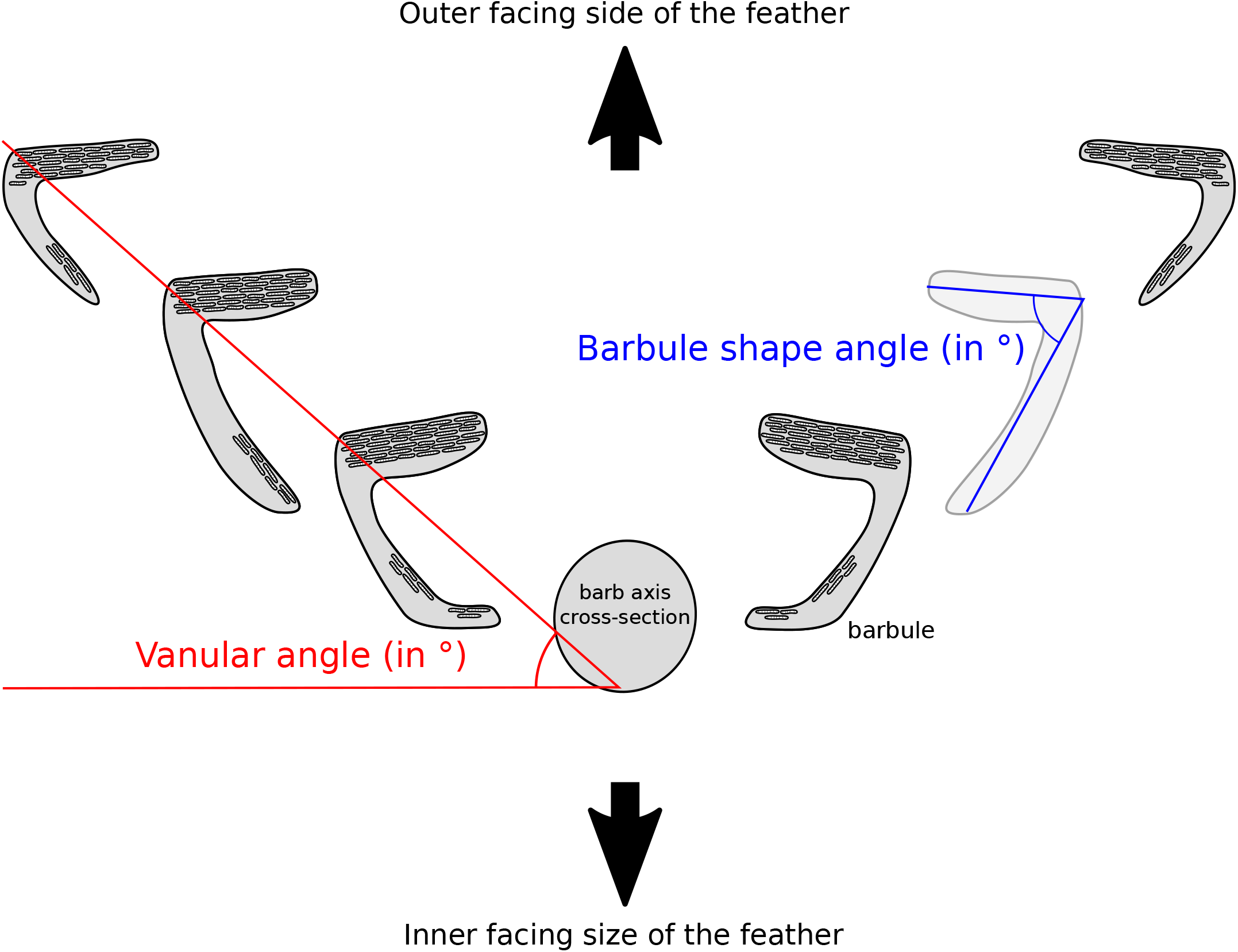
Cross section of a barb and its barbules. Barbule shape angle is displayed in blue and vanular angle in red. Barbules with a large barbule shape angle tend to be flat while barbules with a low barbule shape angle tend to have a sickle shape.

**Supplementary figure 5:**
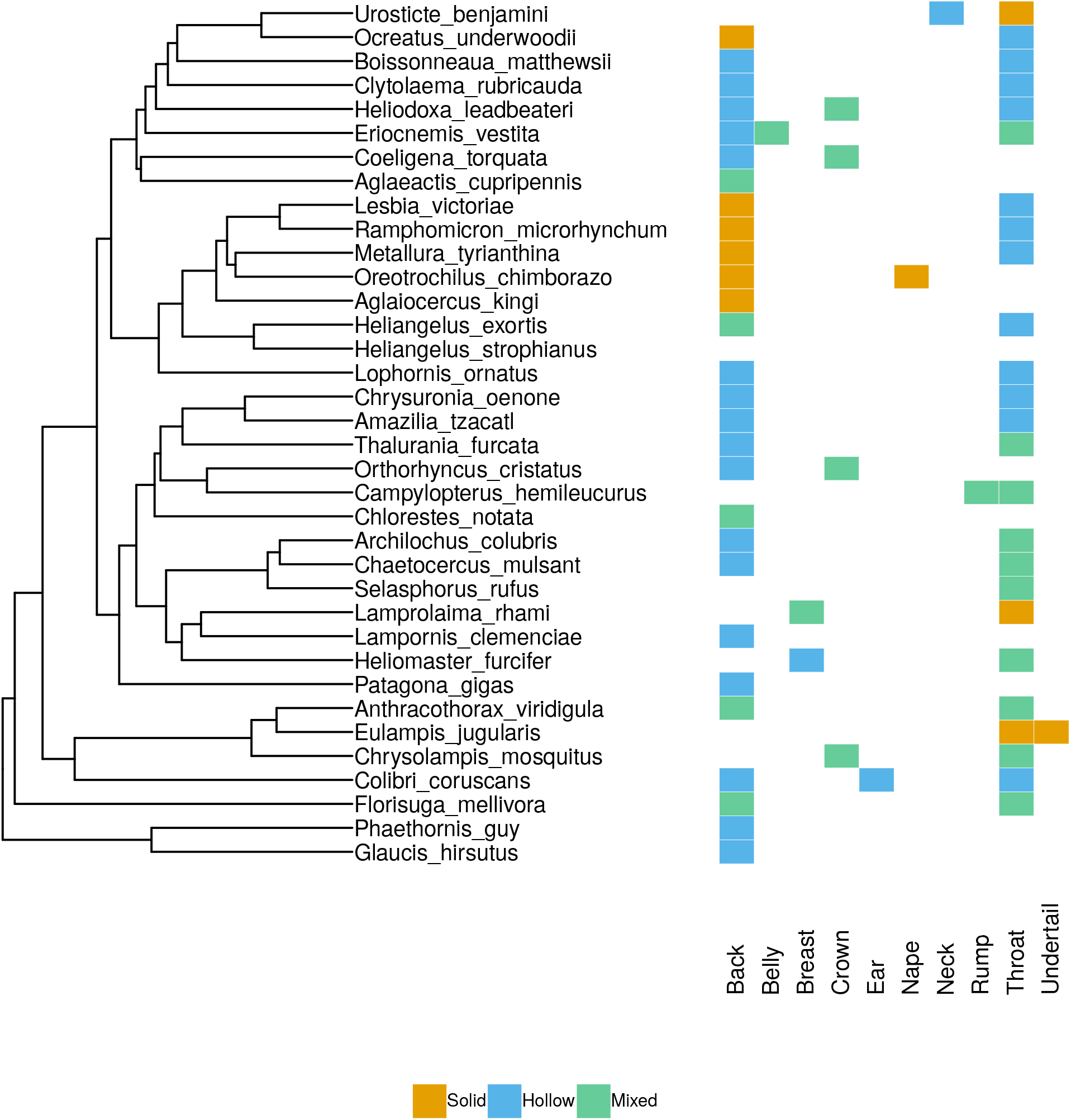
Type of melanosomes on the different patches we measured for each species. The sampling was dependent on which patch was iridescent (and diffuse / directional) for each species. Different patches of the same species can have different types of multilayer structures and patches such as the throat had more often the outer type than patches such as the back.

**Supplementary figure 6:**
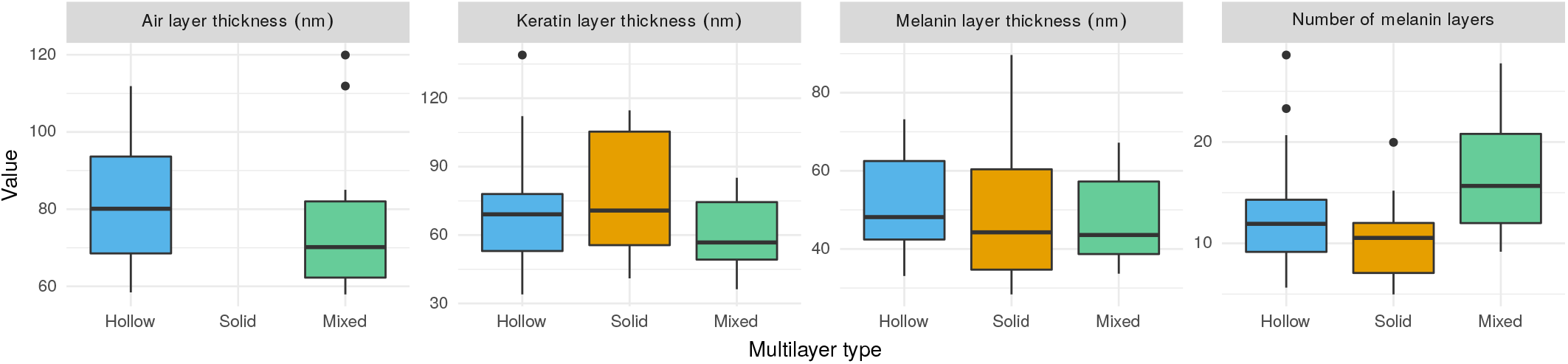
Range of variation of structural parameters for each type of multilayer, estimated on TEM photographs using an automated python script based on OpenCV [23].

**Supplementary figure 7:**
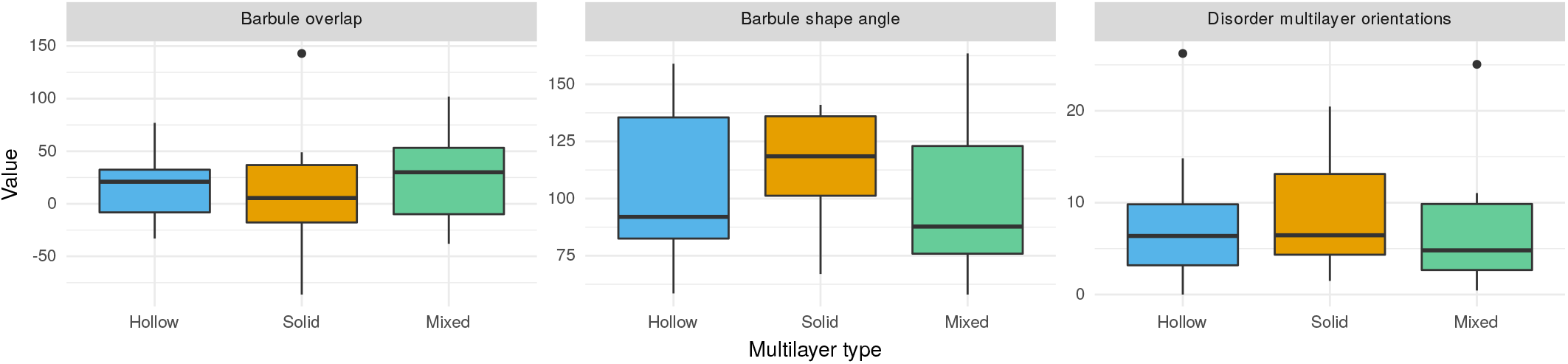
Range of variation of structural parameters for each type of multilayer, estimated on photonic microscopy photographs. The barbule overlap describes how much adjacent barbules overlap and represents how packed and well-organised the barbules are. The barbule shape angle is illustrated in fig. S4. The disorder in the multilayer orientations describe how parallel adjacent multilayers are.

**Supplementary figure 8:**
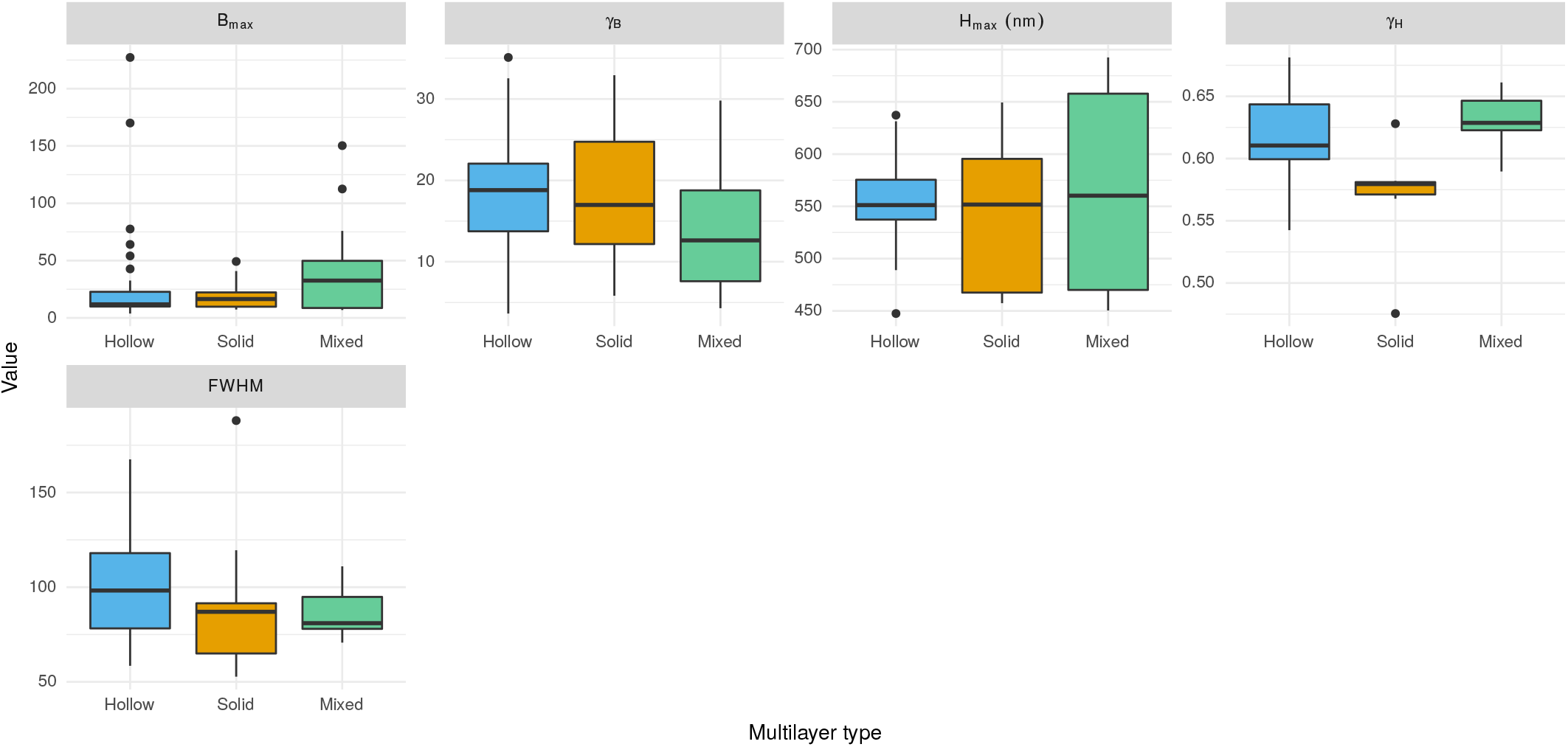
Range of variation of all iridescence parameters for each type of multilayer. The parameters are detailed in Gruson et al. [18]: *B_max_* and *γ_B_* characterise brightness with *B_max_* being the maximum brightness (reached when the illumination coincides with the observer) and *γ_B_* the angle dependency of brightness. Similarly, *H_max_* quantifies the maximum hue (reached when the illumination and the observer and in symmetrical positions relative to the multilayer normal) and *γ_H_* the angle dependency of hue. Finally, FWHM describe the saturation (constant with the angle) with higher values corresponding to less saturated colours.

**Supplementary figure 9:**
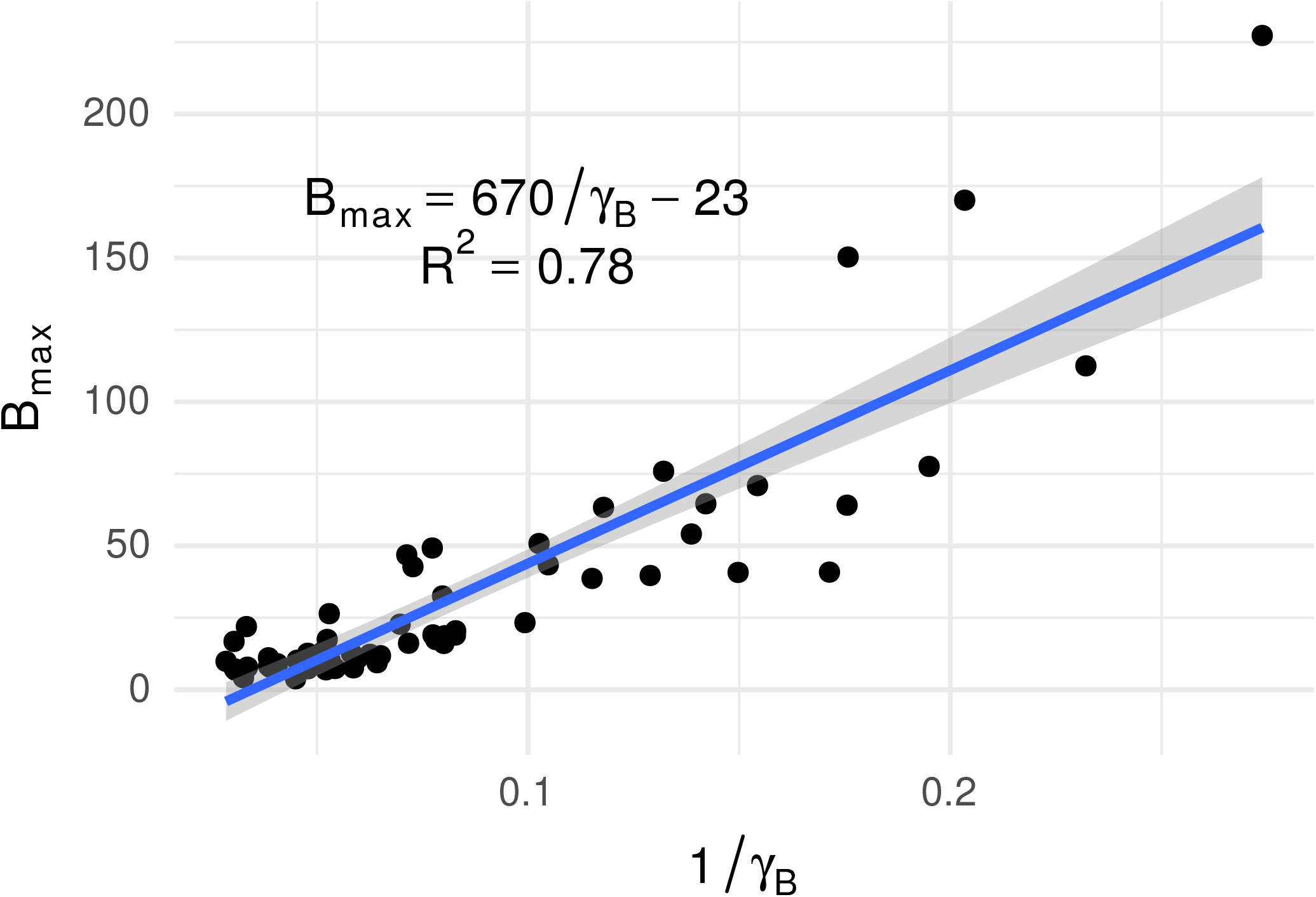
Correlation between *B*_max_ (maximum brightness, reached when the two fibres are in specular position relative to the multilayer structure) and 1*/γ_B_* (directionality *sensu* Osorio and Ham [2]). This is an example of correlation between optical characteristics because of physics (as shown in Gruson et al. [18]).

**Supplementary figure 10:**
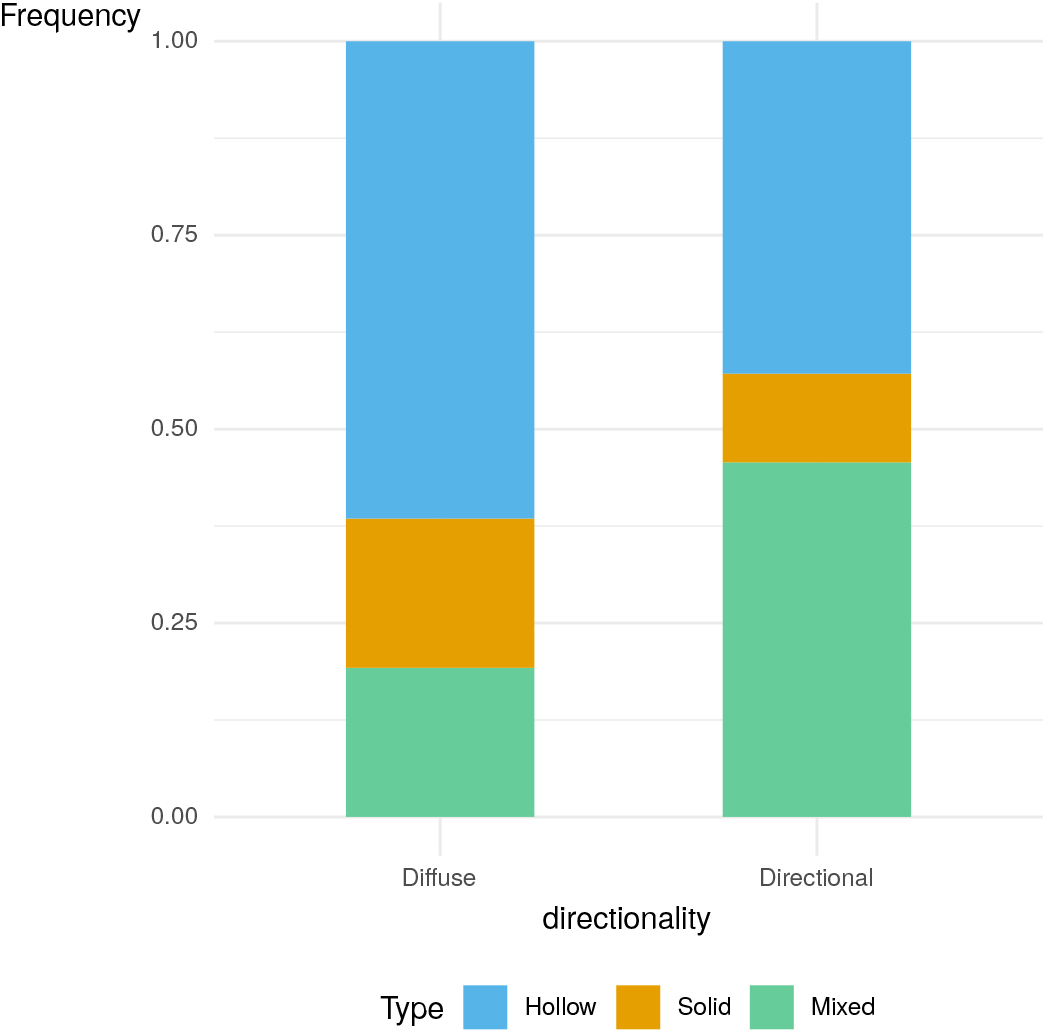
Correlation between multilayer type (hollow, full or mixed) and directionality (diffuse vs directional).

**Supplementary figure 11:**
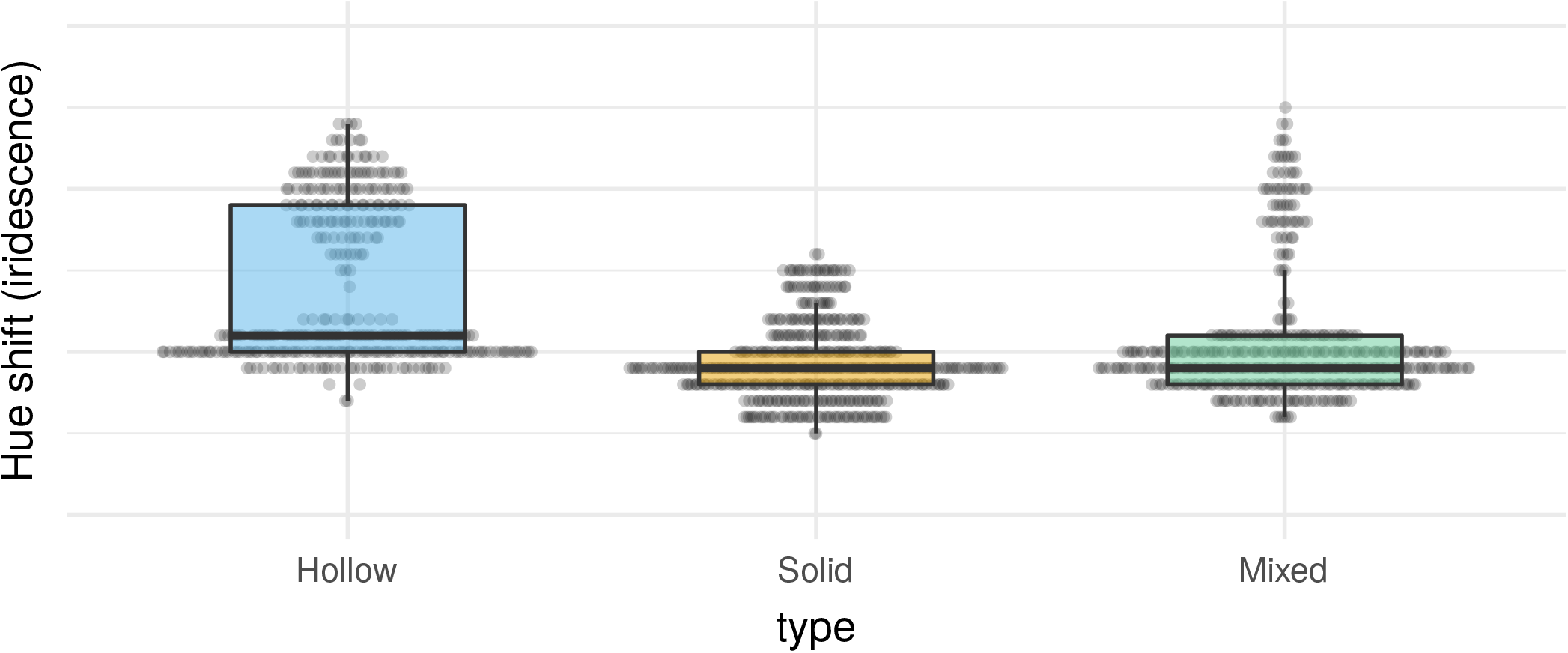
Influence of the multilayer type on the hue shift with the angle change (difference in hue H1 between specular reflection at 0*^◦^* and specular reflection at 10*^◦^*). The data was produced using a transfer matrix model with biologically relevant parameter values.

**Supplementary figure 12:**
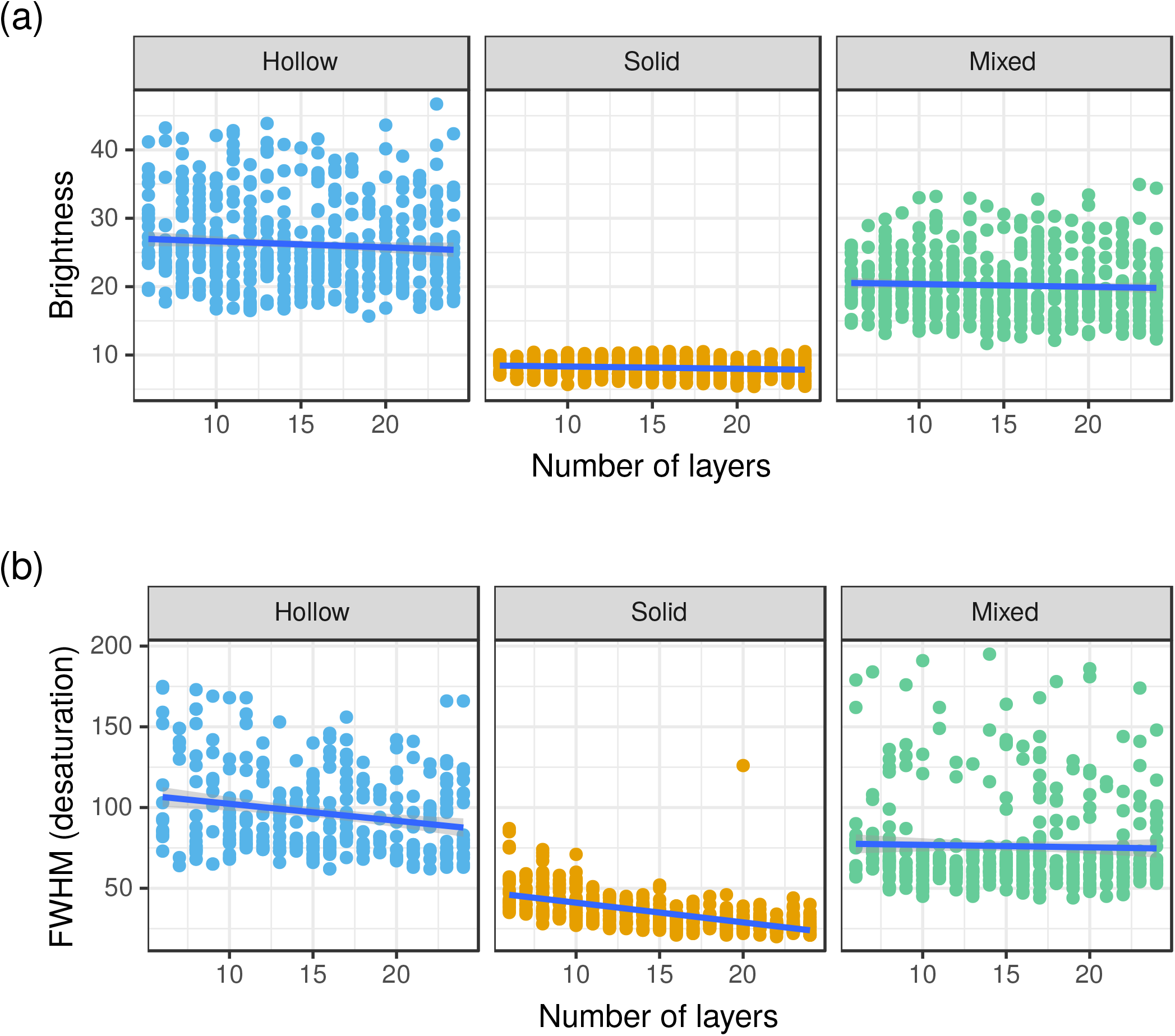
Effect of the number of layers and the type of melanosomes on brightness and FWHM (desaturation). This results from Monte Carlo simulations (500 iterations for each multilayer type) using a transfer-matrix multilayer model. The parameters of each simulation are drawn from a distribution whose range is defined by the analysis of TEM pictures. Statistics analysing the effect of the number of layers and of the multilayer type are presented in tables S4 and S5.

**Supplementary figure 13:**
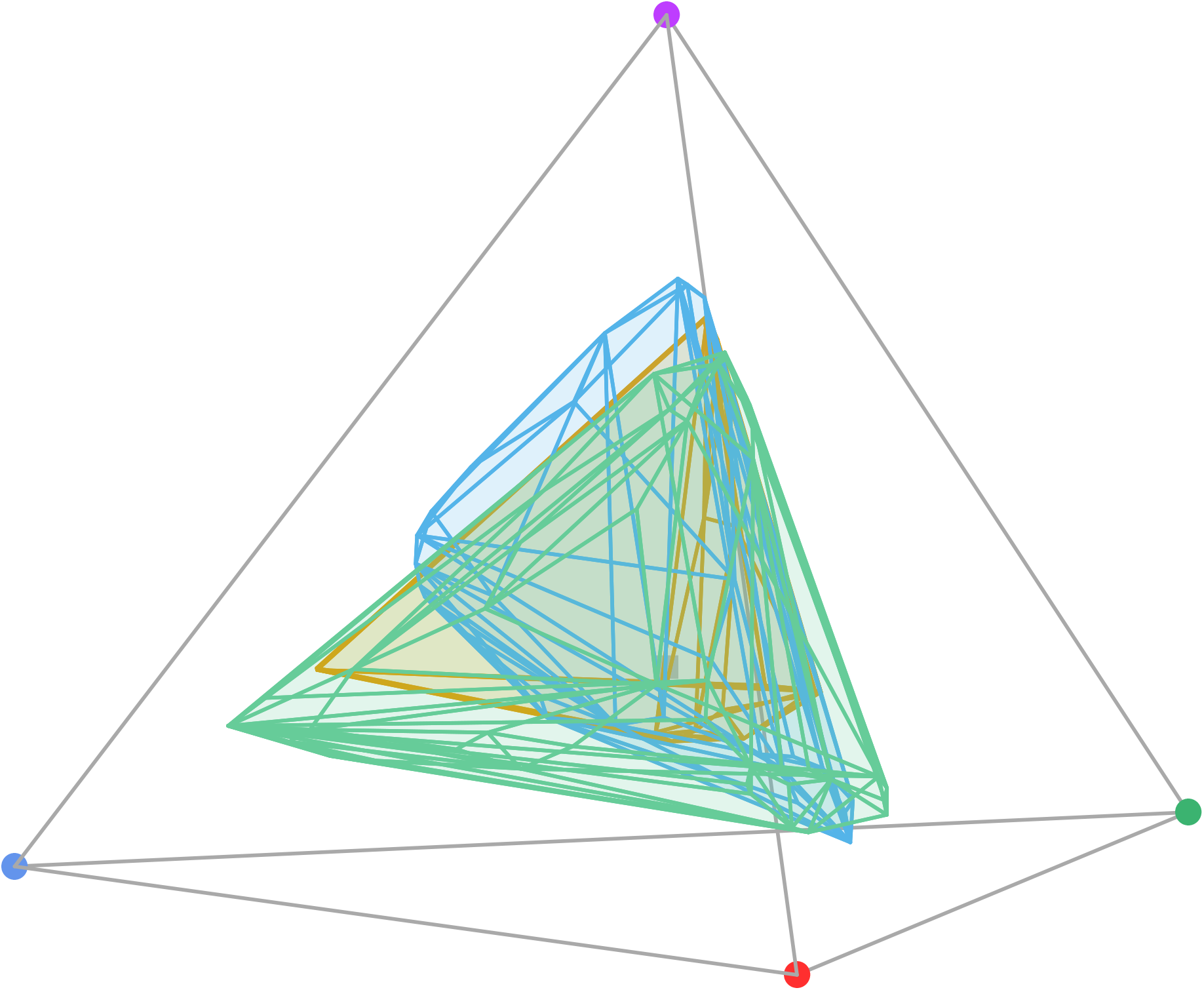
Colour gamut of the different multilayer types. The gamut was determined by computing the convex hull of the set of points obtained by running the result of the transfer matrix simulations in a avian vision model.

**Supplementary table 1:** Iridescence variables (related to the visual signal) and structure characteristics are repeatable within our sample. Repeatability is measured as the intra-class coefficient (ICC) and p-values are estimated by two methods: permutation (p permutation) and likelihood ratio (p likelihood). All repeatability calculations are performed using the rptR R package [57]. Measurement error is also estimated using relative standard deviation (RSD, also called coefficient of variation CV) which compares the standard deviation of several measurements of the same feature to its average.

| Variable | RSD (%) | ICC | p (permutation) | p (likelihood) |
| --- | --- | --- | --- | --- |
| $H_{\max}$ | 0.25 | 1.00 | 0.00 | < 0.0001 |
| $\gamma_H$ | 2.46 | 0.17 | 0.21 | 0.2396 |
| $B_{\max}$ | 16.03 | 0.89 | 0.04 | < 0.0001 |
| $\gamma_B$ | 23.05 | 0.71 | 0.01 | 0.0087 |
| FWHM | 2.27 | 0.66 | 0.00 | < 0.0001 |
| Number of layers | 19.77 | 0.54 | 0.00 | < 0.0001 |
| Melanin layer thickness | 10.79 | 0.54 | 0.00 | < 0.0001 |
| Keratin layer thickness | 15.53 | 0.64 | 0.00 | < 0.0001 |
| Air layer thickness | 8.95 | 0.77 | 0.00 | < 0.0001 |
| Variability in air layer thickness | 14.44 | 0.77 | 0.00 | < 0.0001 |
| Variability in keratin layer thickness | 29.28 | 0.41 | 0.00 | < 0.0001 |
| Variability in melanin layer thickness | 17.37 | 0.45 | 0.00 | < 0.0001 |
| Barbule shape angle | 7.07 | 0.91 | 0.00 | < 0.0001 |
| Barbule overlap | 19.72 | 0.50 | 0.02 | < 0.0001 |

**Supplementary table 2:** Correlation between FWHM (opposite of saturation), overall reflectance 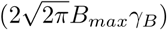, *γ_B_* (inversely related to directionality) and *H_max_* in empirical data from hummingbird feathers.

|  | Estimates(median) | lower.CI(2.5) | upper.CI(97.5) |
| --- | --- | --- | --- |
| (Intercept) | 25.84 | -42.75 | 94.49 |
| totalreflect | -0.01 | -0.02 | 0.00 |
| gammaB | 0.54 | -0.72 | 1.80 |
| Hmax | 0.14 | 0.00 | 0.27 |
| phylogenetic.variance | 2.27 | 0.00 | 264.36 |
| residual.variance | 694.73 | 453.60 | 1114.18 |

**Supplementary table 3:** Correlation between overall reflectance 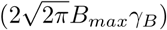, FWHM (opposite of saturation), *γ_B_* (inversely related to directionality) and *H_max_* in empirical data from hummingbird feathers.

|  | Estimates(median) | lower.CI(2.5) | upper.CI(97.5) |
| --- | --- | --- | --- |
| (Intercept) | -559.55 | -2725.62 | 1609.37 |
| FWHM | -7.12 | -16.72 | 2.47 |
| gammaB | -47.01 | -84.28 | -9.77 |
| Hmax | 6.79 | 2.76 | 10.82 |
| phylogenetic.variance | 64.10 | 0.00 | 156309.83 |
| residual.variance | 698401.57 | 463069.31 | 1114969.23 |

**Supplementary table 4:**
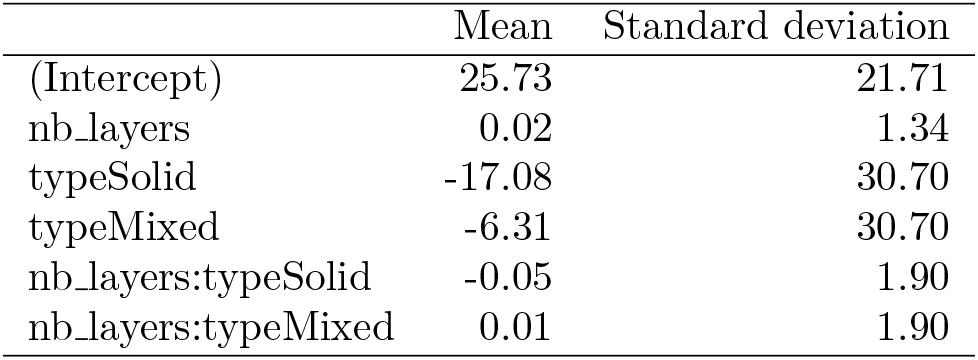
Influence of structural parameters on brightness. Optical theory predicts that brightness (*B*_max_) is controlled by the number of layers and their refractive index (i.e. the type of melanosomes). We test this on simulated data from Monte Carlo transfer matrix simulations using linear models. We find that brightness in simulated data is indeed influenced by the type of melanosomes and by the number of layers. This result is also illustrated in fig. S1.

|  | Mean | Standard deviation |
| --- | --- | --- |
| (Intercept) | 25.73 | 21.71 |
| nb_layers | 0.02 | 1.34 |
| typeSolid | -17.08 | 30.70 |
| typeMixed | -6.31 | 30.70 |
| nb_layers:typeSolid | -0.05 | 1.90 |
| nb_layers:typeMixed | 0.01 | 1.90 |

**Supplementary table 5:** Influence of structural parameters on saturation. Optical theory predicts that FWHM (opposite of saturation) is controlled by the number of layers and their refractive index (i.e. the type of melanosomes), as well as layer thickness. We test this on simulated data from Monte Carlo transfer matrix simulations using linear models. FWHM (opposite of saturation) in simulated data is indeed influenced by the type of melanosomes and by the number of layers for the solid and the mixed type. This result is also illustrated in fig. S2.

|  | Mean | Standard deviation |
| --- | --- | --- |
| (Intercept) | 136.85 | 255.60 |
| nb_layers | -1.21 | 8.56 |
| typeSolid | -48.01 | 242.87 |
| typeMixed | -29.25 | 183.57 |
| melanin_size | 0.24 | 2.14 |
| keratin_size | -0.65 | 1.23 |
| air_size | 0.18 | 2.02 |
| nb_layers:typeSolid | 0.14 | 10.93 |
| nb_layers:typeMixed | 0.83 | 11.36 |

**Supplementary table 6:** Influence of structural parameters on hue. Optical theory predicts that hue (*H*_max_) is controlled by the thickness of each layer and their refractive index (i.e. the type of melanosomes). We test this on simulated data from Monte Carlo transfer matrix simulations using linear models. Hue in the simulated data indeed depends on the type of melanosomes, the thickness of the layers and the interaction of both.

|  | Mean | Standard deviation |
| --- | --- | --- |
| (Intercept) | 5.26 | 402.79 |
| melanin_size | 3.13 | 4.40 |
| keratin_size | 1.64 | 2.51 |
| air_size | 0.84 | 2.78 |
| typeSolid | 48.13 | 425.37 |
| typeMixed | 74.11 | 443.55 |
| interference order (m) | 330.89 | 93.11 |
| melanin_size:typeSolid | -0.57 | 5.55 |
| melanin_size:typeMixed | -0.44 | 5.71 |
| keratin_size:typeSolid | 1.26 | 2.79 |
| keratin_size:typeMixed | -0.17 | 3.13 |
| air_size:typeMixed | -0.38 | 3.88 |

**Supplementary table 7:**
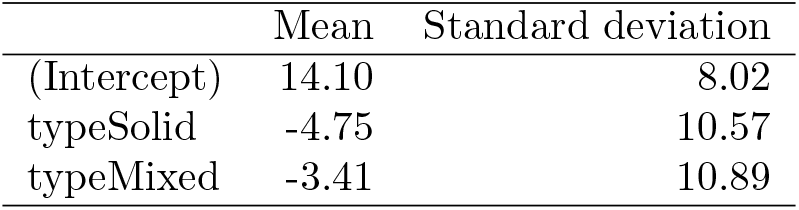
Influence of the multilayer type on hue shift with the change in illumination and observation angle (difference in hue H1 between specular reflection at 0*^◦^* and specular reflection at 10*^◦^*; strongly related to *γ_H_*, as explained in Gruson et al. 18). The linear model was ran on simulated data using a transfer matrix model with biologically relevant parameter values.

**Supplementary table 8:**
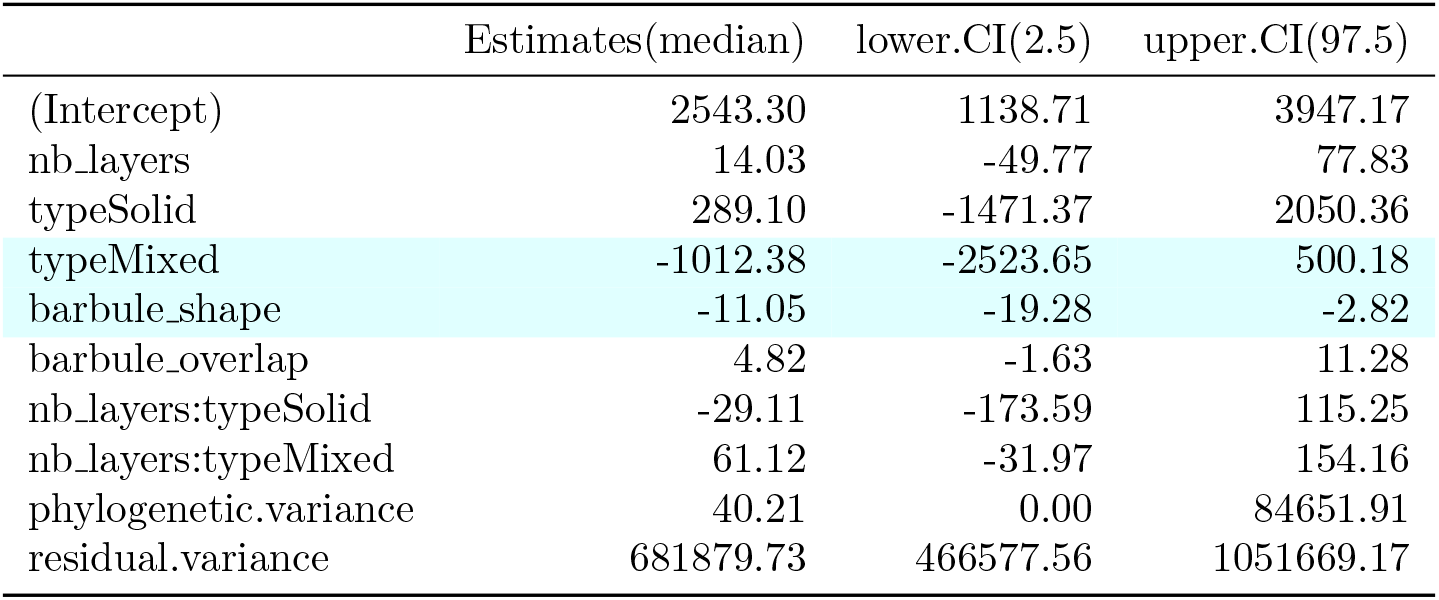
Correlation between total brightness and structural parameters. Optical theory predicts that total brightness (proportional to *B*_max_*γ_B_*) is controlled by the number of layers, their refractive index (i.e. the type of melanosomes) and how packed barbules are (barbule shape and overlap). We test this on empirical data from hummingbird iridescent feathers using `MCMCglmm`. The first column contains explanatory variables, the second one the estimate of the effect size, and the third and fourth one the lower and higher (respectively) bounds of the 95 % credibility interval for the effect size. Significant effects of explanatory variables are shown with a cyan background. This result is also illustrated in fig. 4b.

|  | Estimates(median) | lower.CI(2.5) | upper.CI(97.5) |
| --- | --- | --- | --- |
| (Intercept) | 2543.30 | 1138.71 | 3947.17 |
| nb_layers | 14.03 | -49.77 | 77.83 |
| typeSolid | 289.10 | -1471.37 | 2050.36 |
| typeMixed | -1012.38 | -2523.65 | 500.18 |
| barbule_shape | -11.05 | -19.28 | -2.82 |
| barbule_overlap | 4.82 | -1.63 | 11.28 |
| nb_layers:typeSolid | -29.11 | -173.59 | 115.25 |
| nb_layers:typeMixed | 61.12 | -31.97 | 154.16 |
| phylogenetic.variance | 40.21 | 0.00 | 84651.91 |
| residual.variance | 681879.73 | 466577.56 | 1051669.17 |

**Supplementary table 9:** Correlation between directionality and structural parameters. Optical theory predicts that angular dependency of brightness *γ_B_* (inversely proportional to directionality *sensu* Osorio and Ham [2]) is controlled by how well-arranged barbules are (barbule shape, overlap and alignment). We test this on empirical data from hummingbird iridescent feathers using `MCMCglmm`. The first column contains explanatory variables, the second one the estimate of the effect size, and the third and fourth one the lower and higher (respectively) bounds of the 95 % credibility interval for the effect size. Significant effects of explanatory variables are shown with a cyan background. This result is also illustrated in fig. 4a.

|  | Estimates(median) | lower.CI(2.5) | upper.CI(97.5) |
| --- | --- | --- | --- |
| (Intercept) | 3.76 | -3.72 | 11.13 |
| barbule_shape | 0.12 | 0.05 | 0.19 |
| barbule_overlap | -0.05 | -0.10 | 0.01 |
| align_disorder | 0.17 | -0.21 | 0.55 |
| phylogenetic.variance | 0.52 | 0.00 | 16.14 |
| residual.variance | 50.48 | 34.54 | 76.92 |

**Supplementary table 10:** Correlation between hue (*H_max_*) and structural parameters. Optical theory predicts that hue *H*_max_ is controlled by layer thickness and refractive index (i.e. multilayer type). We test this on empirical data from hummingbird iridescent feathers using `MCMCglmm`. The first column contains explanatory variables, the second one the estimate of the effect size, and the third and fourth one the lower and higher (respectively) bounds of the 95 % credibility interval for the effect size. Significant effects of explanatory variables are shown with a cyan background.

|  | Estimates(median) | lower.CI(2.5) | upper.CI(97.5) |
| --- | --- | --- | --- |
| (Intercept) | 530.52 | 434.10 | 628.52 |
| melanin_thickness | -3.95 | -6.91 | -1.03 |
| keratin_thickness | 1.31 | -0.15 | 2.82 |
| air_thickness | 1.07 | -0.27 | 2.39 |
| typeSolid | 51.99 | -83.20 | 189.78 |
| typeMixed | -48.62 | -176.52 | 83.84 |
| interference order (m) | 93.95 | 67.99 | 119.55 |
| melanin_thickness:typeSolid | 1.92 | -2.01 | 5.79 |
| melanin_thickness:typeMixed | 9.64 | 3.69 | 15.85 |
| keratin_thickness:typeSolid | -1.09 | -3.02 | 0.87 |
| keratin_thickness:typeMixed | -6.06 | -9.56 | -2.87 |
| air_thickness:typeMixed | -0.39 | -2.32 | 1.55 |
| phylogenetic.variance | 177.48 | 0.01 | 2266.77 |
| residual.variance | 973.09 | 297.37 | 1893.45 |

**Supplementary table 11:** Correlation between FWHM (opposite of saturation) and structural parameters. Optical theory predicts that saturation is controlled by the variance in layer thickness, the number of layers and their refractive index (i.e. multilayer type) as well as disorder in the alignment of the multilayers. We test this on empirical data from hummingbird iridescent feathers using `MCMCglmm`. The first column contains explanatory variables, the second one the estimate of the effect size, and the third and fourth one the lower and higher (respectively) bounds of the 95 % credibility interval for the effect size. Significant effects of explanatory variables are shown with a cyan background.

|  | Estimates(median) | lower.CI(2.5) | upper.CI(97.5) |
| --- | --- | --- | --- |
| (Intercept) | 83.79 | 39.06 | 128.68 |
| mel_varsize | -0.56 | -2.27 | 1.18 |
| ker_varsize | -0.01 | -1.09 | 1.13 |
| air_varsize | 0.45 | -0.93 | 1.80 |
| typeSolid | -54.15 | -109.99 | -0.41 |
| typeMixed | -37.85 | -84.53 | 8.81 |
| nb_layers | -1.04 | -2.73 | 0.65 |
| align_disorder | 1.20 | -0.05 | 2.45 |
| barbule_shape | 0.25 | 0.00 | 0.50 |
| typeSolid:nb_layers | 3.28 | -1.80 | 8.53 |
| typeMixed:nb_layers | 1.76 | -0.95 | 4.48 |
| phylogenetic.variance | 88.77 | 0.01 | 473.16 |
| residual.variance | 348.87 | 176.42 | 658.59 |

**Supplementary table 12:** Correlation between melanin and keratin layer thicknesses using `MCMCglmm`. The first column contains explanatory variables, the second one the estimate of the effect size, and the third and fourth one the lower and higher (respectively) bounds of the 95 % credibility interval for the effect size. Significant effects of explanatory variables are shown with a cyan background.

|  | Estimates(median) | lower.CI(2.5) | upper.CI(97.5) |
| --- | --- | --- | --- |
| (Intercept) | 15.74 | 6.72 | 23.40 |
| keratin_size | 0.51 | 0.41 | 0.62 |
| phylogenetic.variance | 5.41 | 0.00 | 70.55 |
| residual.variance | 55.64 | 30.58 | 87.55 |

**Supplementary table 13:** Correlation between melanin and air layer thicknesses using `MCMCglmm`. The first column contains explanatory variables, the second one the estimate of the effect size, and the third and fourth one the lower and higher (respectively) bounds of the 95 % credibility interval for the effect size. Significant effects of explanatory variables are shown with a cyan background.

|  | Estimates(median) | lower.CI(2.5) | upper.CI(97.5) |
| --- | --- | --- | --- |
| (Intercept) | 18.21 | 2.88 | 34.16 |
| air_thickness | 0.41 | 0.22 | 0.60 |
| phylogenetic.variance | 2.92 | 0.00 | 86.14 |
| residual.variance | 92.40 | 53.90 | 146.26 |

**Supplementary table 14:** Correlation between keratin and air layer thicknesses using `MCMCglmm`. The first column contains explanatory variables, the second one the estimate of the effect size, and the third and fourth one the lower and higher (respectively) bounds of the 95 % credibility interval for the effect size. Significant effects of explanatory variables are shown with a cyan background.

|  | Estimates(median) | lower.CI(2.5) | upper.CI(97.5) |
| --- | --- | --- | --- |
| (Intercept) | 32.58 | 3.19 | 62.41 |
| air_thickness | 0.45 | 0.11 | 0.78 |
| phylogenetic.variance | 128.86 | 0.04 | 416.43 |
| residual.variance | 242.14 | 137.09 | 448.51 |

## References

1. S. M. Doucet and M. G. Meadows. Iridescence: A functional perspective. Journal of The Royal Society Interface, 6(Suppl 2):S115–S132, Apr. 2009. doi: 10.1098/rsif.2008.0395.focus.

2. D. C. Osorio and A. D. Ham. Spectral reflectance and directional properties of structural coloration in bird plumage. Journal of Experimental Biology, 205(14):2017–2027, July 2002.

3. H. Gruson, M. Elias, J. L. Parra, C. Andraud, S. Berthier, C. Doutrelant, and D. Gomez. Distribution of iridescent colours in hummingbird communities results from the interplay between selection for camouflage and communication. bioRxiv, page 586362, Apr. 2019. doi: 10.1101/586362.

4. J. Del Hoyo, A. Elliott, J. Sargatal, D. A. Christie, and E. de Juana. Handbook of the Birds of the World Alive. hbw.com, 2017.

5. J. Dorst. Recherches sur la structure des plumes des trochilidés. PhD thesis, Université de Paris, Paris, 1951. OCLC: 14220401.

6. L. D’Alba and M. D. Shawkey. Melanosomes: Biogenesis, Properties, and Evolution of an Ancient Organelle. Physiological Reviews, 99(1):1–19, Sept. 2018. doi: 10.1152/physrev.00059.2017.

7. [7] H. Dürrer. Schillerfarben der Vogelfeder als Evolutionsproblem. PhD thesis, Medizinischen Fakultät der Universität Basel, 1975.

8. C. H. Greenewalt, W. Brandt, and D. D. Friel. Iridescent colors of hummingbird feathers. Journal of the Optical Society of America, 50(10):1005–1013, Oct. 1960. doi: 10.1364/JOSA.50.001005.

9. W. J. Schmidt and H. Ruska. Über das schillernde Federmelanin bei *Heliangelus* und *Lophophorus*. Zeitschrift für Zellforschung und Mikroskopische Anatomie, 57(1), 1962. doi: 10.1007/BF00338926.

10. M. D. Shawkey, N. I. Morehouse, and P. Vukusic. A protean palette: Colour materials and mixing in birds and butterflies. Journal of The Royal Society Interface, 6(Suppl 2):S221–S231, 2009. doi: 10.1098/rsif.2008.0459.focus.

11. M. A. Giraldo, J. L. Parra, and D. G. Stavenga. Iridescent colouration of male Anna’s hummingbird (*Calypte anna*) caused by multilayered barbules. Journal of Comparative Physiology A, Oct. 2018. doi: 10.1007/s00359-018-1295-8.

12. K. K. Nordén, J. W. Faber, F. Babarović, T. L. Stubbs, T. Selly, J. D. Schiffbauer, P. P. Štefanić, G. Mayr, F. M. Smithwick, and J. Vinther. Melanosome diversity and convergence in the evolution of iridescent avian feathers—Implications for paleocolor reconstruction. Evolution, 73(1):15–27, 2019. doi: 10.1111/evo.13641.

13. R. Maia, D. R. Rubenstein, and M. D. Shawkey. Key ornamental innovations facilitate diversification in an avian radiation. Proceedings of the National Academy of Sciences, 110(26):10687–10692, June 2013. doi: 10.1073/pnas.1220784110.

14. G. E. Hill. Female house finches prefer colourful males: Sexual selection for a condition-dependent trait. Animal Behaviour, 40(3):563–572, Sept. 1990. doi: 10.1016/S0003-3472(05)80537-8.

15. A. Loyau, D. Gomez, B. Moureau, M. Théry, N. S. Hart, M. S. Jalme, A. T. D. Bennett, and G. Sorci. Iridescent structurally based coloration of eyespots correlates with mating success in the peacock. Behavioral Ecology, 18(6): 1123–1131, Nov. 2007. doi: 10.1093/beheco/arm088.

16. D. J. Kemp. Female butterflies prefer males bearing bright iridescent ornamentation. Proceedings of the Royal Society of London B: Biological Sciences, 274(1613):1043–1047, Apr. 2007. doi: 10.1098/rspb.2006.0043.

17. D. J. Kemp. Female mating biases for bright ultraviolet iridescence in the butterfly *Eurema hecabe* (Pieridae). Behavioral Ecology, 19(1):1–8, Jan. 2008. doi: 10.1093/beheco/arm094.

18. H. Gruson, C. Andraud, W. Daney de Marcillac, S. Berthier, M. Elias, and D. Gomez. Quantitative characterization of iridescent colours in biological studies: A novel method using optical theory. Interface Focus, 9(1):20180049, Feb. 2019. doi: 10.1098/rsfs.2018.0049.

19. J. Felsenstein. Phylogenies and the comparative method. The American Naturalist, 125(1):1–15, Jan. 1985. doi: 10.1086/284325.

20. W. Jetz, G. H. Thomas, J. B. Joy, K. Hartmann, and A. O. Mooers. The global diversity of birds in space and time. Nature, 491(7424):444–448, Nov. 2012. doi: 10.1038/nature11631.

21. P.-C. Bürkner. Brms: An R package for Bayesian multilevel models using Stan. Journal of Statistical Software, 80 (1):1–28, 2017. doi: 10.18637/jss.v080.i01.

22. R Core Team. R: A Language and Environment for Statistical Computing, 2017.

23. Opencv: Open Source Computer Vision Library. OpenCV, 2017.

24. Python Software Foundation. Python Language Reference.

25. L. Bolla. EMpy - ElectroMagnetic Python, 2017.

26. P. Yeh. Optical Waves in Layered Media. Wiley Series in Pure and Applied Optics. Wiley-Interscience, Hoboken, NJ, 2005. ISBN 978-0-471-73192-4. OCLC: 255155115.

27. M. C. Stoddard and R. O. Prum. Evolution of avian plumage color in a tetrahedral color space: A phylogenetic analysis of New World buntings. The American Naturalist, 171(6):755–776, June 2008. doi: 10.1086/587526.

28. R. Maia, C. M. Eliason, P.-P. Bitton, S. M. Doucet, and M. D. Shawkey. Pavo: An R package for the analysis, visualization and organization of spectral data. Methods in Ecology and Evolution, 4(10):906–913, Oct. 2013. doi: 10.1111/2041-210X.12069.

29. J. D. Hadfield. MCMC Methods for Multi-Response Generalized Linear Mixed Models: The MCMCglmm R Package. Journal of Statistical Software, Feb. 2010. doi: 10.18637/jss.v033.i02.

30. R Core Team. R: A Language and Environment for Statistical Computing, 2018.

31. K. Delhey, M. Hall, S. A. Kingma, and A. Peters. Increased conspicuousness can explain the match between visual sensitivities and blue plumage colours in fairy-wrens. Proceedings of the Royal Society B: Biological Sciences, 280 (1750):20121771, Jan. 2013. doi: 10.1098/rspb.2012.1771.

32. M. Pagel and F. Lutzoni. Accounting for phylogenetic uncertainty in comparative studies of evolution and adaptation. In M. Lässig and A. Valleriani, editors, Biological Evolution and Statistical Physics, volume 585 of *Lecture Notes in Physics*. Springer Berlin Heidelberg, Berlin, Heidelberg, 2002. ISBN 978-3-540-43188-6. doi: 10.1007/3-540-45692-9.

33. P. de Villemereuil, J. A. Wells, R. D. Edwards, and S. P. Blomberg. Bayesian models for comparative analysis integrating phylogenetic uncertainty. BMC Evolutionary Biology, 12(1):102, June 2012. doi: 10.1186/1471-2148-12-102.

34. T. Guillerme and K. Healy. mulTree: A package for running MCMCglmm analysis on multiple trees. Zenodo, Nov. 2014.

35. R. Borges, J. P. Machado, C. Gomes, A. P. Rocha, and A. Antunes. Measuring phylogenetic signal between categorical traits and phylogenies. Bioinformatics, 35(11):1862–1869, June 2019. doi: 10.1093/bioinformatics/bty800.

36. M. A. Rodŕıguez-Gironés and L. Santamaŕıa. Why are so many bird flowers red? PLOS Biology, 2(10):e350, Oct. 2004. doi: 10.1371/journal.pbio.0020350.

37. H. L. Leertouwer, B. D. Wilts, and D. G. Stavenga. Refractive index and dispersion of butterfly chitin and bird keratin measured by polarizing interference microscopy. Optics Express, 19(24):24061–24066, Nov. 2011. doi: 10.1364/OE.19.024061.

38. D. G. Stavenga, H. L. Leertouwer, D. C. Osorio, and B. D. Wilts. High refractive index of melanin in shiny occipital feathers of a bird of paradise. Light: Science & Applications, 4(1):e243, Jan. 2015. doi: 10.1038/lsa.2015.16.

39. K. Kjernsmo, J. R. Hall, C. Doyle, N. Khuzayim, I. C. Cuthill, N. E. Scott-Samuel, and H. M. Whitney. Iridescence impairs object recognition in bumblebees. Scientific Reports, 8(1):8095, May 2018. doi: 10.1038/s41598-018-26571-6.

40. R. O. Prum. Anatomy, physics, and evolution of structural colors. In G. E. Hill and K. J. McGraw, editors, Bird Coloration, Volume 1: Mechanisms and Measurements, volume 1 of Bird Coloration, page 640. Harvard University Press, Feb. 2006. ISBN 978-0-674-01893-8.

41. C. M. Eliason, P.-P. Bitton, and M. D. Shawkey. How hollow melanosomes affect iridescent colour production in birds. Proc. R. Soc. B, 280(1767):20131505, Sept. 2013. doi: 10.1098/rspb.2013.1505.

42. J. A. McGuire, C. C. Witt, J. V. J. Remsen, A. Corl, D. L. Rabosky, D. L. Altshuler, and R. Dudley. Molecular phylogenetics and the diversification of hummingbirds. Current Biology, 24(8):910–916, Apr. 2014. doi: 10.1016/j.cub.2014.03.016.

43. M. F. Land. A multilayer interference reflector in the eye of the scallop, *Pecten maximus*. Journal of Experimental Biology, 45(3):433–447, Dec. 1966.

44. D. J. Brink and M. E. Lee. Thin-film biological reflectors: Optical characterization of the *Chrysiridia croesus* moth. Applied Optics, 37(19):4213–4217, July 1998. doi: 10.1364/AO.37.004213.

45. J. Chae and S. Nishida. Spectral patterns of the iridescence in the males of *Sapphirina* (Copepoda: Poecilostomatoida). Journal of the Marine Biological Association of the United Kingdom, 79(3):437–443, June 1999. doi: 10.1017/S0025315498000563.

46. D. G. Stavenga, B. D. Wilts, H. L. Leertouwer, and T. Hariyama. Polarized iridescence of the multilayered elytra of the Japanese jewel beetle, *Chrysochroa fulgidissima*. Philosophical Transactions of the Royal Society B: Biological Sciences, 366(1565):709–723, Mar. 2011. doi: 10.1098/rstb.2010.0197.

47. C. M. Eliason, R. Maia, and M. D. Shawkey. Modular color evolution facilitated by a complex nanostructure in birds. Evolution, 69(2):357–367, Feb. 2015. doi: 10.1111/evo.12575.

48. M. F. Land. The physics and biology of animal reflectors. Progress in Biophysics and Molecular Biology, 24:75–106, Jan. 1972. doi: 10.1016/0079-6107(72)90004-1.

49. S. Kinoshita, S. Yoshioka, and J. Miyazaki. Physics of structural colors. Reports on Progress in Physics, 71(7): 076401, 2008. doi: 10.1088/0034-4885/71/7/076401.

50. D. G. Stavenga, H. L. Leertouwer, and B. D. Wilts. Magnificent magpie colours by feathers with layers of hollow melanosomes. Journal of Experimental Biology, 221(4):jeb174656, Feb. 2018. doi: 10.1242/jeb.174656.

51. S. Berthier, E. Charron, and J. Boulenguez. Morphological structure and optical properties of the wings of Morphidae. Insect Science, 13(2):145–158, 2006. doi: 10.1111/j.1744-7917.2006.00077.x.

52. D. G. Stavenga, H. L. Leertouwer, N. J. Marshall, and D. C. Osorio. Dramatic colour changes in a bird of paradise caused by uniquely structured breast feather barbules. Proceedings of the Royal Society of London B: Biological Sciences, 278(1715):2098–2104, July 2011. doi: 10.1098/rspb.2010.2293.

53. B. D. Wilts, K. Michielsen, H. D. Raedt, and D. G. Stavenga. Sparkling feather reflections of a bird-of-paradise explained by finite-difference time-domain modeling. Proceedings of the National Academy of Sciences, 111(12): 4363–4368, Mar. 2014. doi: 10.1073/pnas.1323611111.

54. H. Dürrer and W. Villiger. Schillerradien des Goldkuckucks (Chrysococcyx cupreus (Shaw)) im Elektronenmikroskop. Zeitschrift für Zellforschung und Mikroskopische Anatomie, 109(3):407–413, Sept. 1970. doi: 10.1007/BF02226912.

55. H. Sick. Morphologisch-funktionelle Untersuchungen über die Feinstruktur der Vogelfeder. Journal für Ornithologie, 85(2):206–372, Apr. 1937. doi: 10.1007/BF01905702.

56. C. H. Greenewalt. Hummingbirds. Dover Publications, New York, Feb. 1991. ISBN 978-0-486-26431-8.

57. S. Nakagawa and H. Schielzeth. Repeatability for Gaussian and non-Gaussian data: A practical guide for biologists. Biological Reviews, 85(4):935–956, Nov. 2010. doi: 10.1111/j.1469-185X.2010.00141.x.

